# mitoBK_Ca_ is functionally expressed in murine and human breast cancer cells and potentially contributes to metabolic reprogramming

**DOI:** 10.1101/2023.10.02.560571

**Authors:** Helmut Bischof, Selina Maier, Piotr Koprowski, Bogusz Kulawiak, Sandra Burgstaller, Joanna Jasińska, Kristian Serafimov, Monika Zochowska, Dominic Gross, Werner Schroth, Lucas Matt, David Arturo Juarez Lopez, Ying Zhang, Irina Bonzheim, Florian A. Büttner, Falko Fend, Matthias Schwab, Andreas L. Birkenfeld, Roland Malli, Michael Lämmerhofer, Piotr Bednarczyk, Adam Szewczyk, Robert Lukowski

## Abstract

Alterations in the function of K^+^ channels such as the voltage- and Ca^2+^ activated K^+^ channel of large conductance (BK_Ca_) reportedly promote breast cancer (BC) development and progression. Underlying molecular mechanisms remain, however, elusive. Here, we provide electrophysiological evidence for a BK_Ca_ splice variant localized to the inner mitochondrial membrane of murine and human BC cells (mitoBK_Ca_). Through a combination of genetic knockdown and knockout along with cell permeable BK_Ca_ channel blocker, we show that mitoBK_Ca_ modulates overall cellular and mitochondrial energy production and mediates the metabolic rewiring referred to as the “Warburg effect”, thereby promoting BC cell proliferation in the presence and absence of oxygen. Additionally, we detect mitoBK_Ca_ and BK_Ca_ transcripts in low or high abundance, respectively, in clinical BC specimens. Together, our results emphasize, that targeting mitoBK_Ca_ could represent a treatment strategy for selected BC patients in future.

## Introduction

Cancer represents a complex disease characterized by unconstrained cell proliferation and the spread of malignant cells in the body (Kalia, 2015; Seyfried and Shelton, 2010). It is one of the leading causes of death worldwide, with millions of new cases diagnosed each year (Sung et al., 2021). Globally, the most prevalent form of cancer represents breast cancer (BC) (Sung et al., 2021). Despite many available anti-cancer treatments which largely depend on the steroid and epidermal growth factor (HER2) receptor status (Dunnwald et al., 2007), cancer cells frequently escape from existing therapies due to adaptions (P. Wu et al., 2021). Therefore, the identification of novel targets and therapeutic strategies that confer benefits for at least a subset of patients, whose cancer displays specific molecular or cellular features, is of utmost relevance.

Important factors that emerged as new cancer targets are ion channels (Li and Xiong, 2011). Especially alterations in the expression levels and function of potassium ion (K^+^) channels are critically related to cancer malignancy and progression (Li et al., 2023, p. 0). One of these channels represents the calcium ion (Ca^2+^) and voltage-activated K^+^ channel of large conductance (BK_Ca_) (Mohr et al., 2022). Canonical BK_Ca_ channels usually localize in the plasma membrane (PM) of cells, and contribute to the regulation of the cytosolic K^+^ content, the PM potential (ΔΨ_PM_), cell cycle, and cell motility, as well as regulatory volume, changes (Burgstaller et al., 2022a). Opening of BK_Ca_ channels results in K^+^ efflux, increasing the electrochemical driving force for Ca^2+^ entry into the cancer cell and affecting pathological cell growth and death (Ouadid-Ahidouch and Ahidouch, 2013). Accordingly, in BC cells (BCCs), an upregulation of BK_Ca_ has been associated with increased malignancy (Huang and Jan, 2014; Mohr et al., 2022; Oeggerli et al., 2012). However, besides their localization in the PM, several K^+^ channels, including BK_Ca_, have also been identified in the inner mitochondrial membrane (IMM) (mitoBK_Ca_), a topic which has been extensively reviewed recently (Checchetto et al., 2021; Kulawiak and Szewczyk, 2022; Szabo and Szewczyk, 2023; Szewczyk et al., 2009; Wrzosek et al., 2020). mitoBK_Ca_ was first described by Siemen et al. in a human glioma cell line more than 20 years ago (Siemen et al., 1999). So far, mitoBK_Ca_ has further been found e.g. in bronchial epithelial cells (Dabrowska et al., 2022), neurons (Douglas et al., 2006), skeletal muscle cells (Skalska et al., 2008), and in cardiac myocytes (Xu et al., 2002). In the latter, a splice variant, the BK_Ca_-DEC isoform containing a unique C-terminal exon of 50 amino acids forms the functional mitoBK_Ca_ channel at the IMM (Singh et al., 2013). Little, however, is known about the molecular identity of mitoBK_Ca_ in other cell types, and mitochondrial localization of BK_Ca_ in BCCs has not been demonstrated so far.

Cancer cells show increased energy demands due to their high proliferation rates. Thus, tumor cells compensate for their elevated energy demand by increasing metabolic activities and adapting to nutrient-poor metabolic niches in the tumor microenvironment, allowing them to overcome oxygen (O_2_)-dependent mitochondrial metabolism (Eales et al., 2016; Gross et al., 2022; Jang et al., 2013; Nazemi and Rainero, 2020). This metabolic switch from oxidative phosphorylation to glycolysis, frequently referred to as the “Warburg effect”, describes the phenomenon that cancer cells rather secrete lactate to the extracellular matrix (ECM), instead of utilizing pyruvate to fuel the TCA cycle (Warburg, 1924). This lactate secretion towards the ECM was shown to promote multiple microenvironmental cues causing tumor progression (de la Cruz-López et al., 2019). Interestingly, extracellular K^+^ may also impair effector T-cell function, and, thus, the anti-tumor immune response (Eil et al., 2016), while, within the cancer cell, functions of several glycolytic enzymes rely on K^+^ (Bischof et al., 2021; Gohara and Di Cera, 2016). Further, the presence of mitoBK_Ca_ contributes to the K^+^ entry into the mitochondrial matrix to interfere with mitochondrial volume changes and the mitochondrial membrane potential (ΔΨ_mito_) (Krabbendam et al., 2018). Consequently, (mito)/BK_Ca_-dependent mechanisms may severely affect energy production pathways and the resulting energy supply of cancer cells.

To elucidate how BK_Ca_ contributes to increasing BCC malignancy (Oeggerli et al., 2012), we utilized BK_Ca_ pro- and deficient human BCC lines including MDA-MB-453 and MCF-7 cells, and primary murine BCCs derived from the mouse mammary tumor polyoma middle T-antigen (MMTV-PyMT)-induced wild type (WT) or BK_Ca_ knock-out (BK-KO) BC model (Mohr et al., 2022). In these cells, either isoforms of BK_Ca_ were expressed, or BK_Ca_ was pharmacologically blocked by paxilline or iberiotoxin, two frequently used inhibitors of BK_Ca_ that either penetrate the cell or act exclusively at PM localized channels, respectively (Candia et al., 1992; Zhou and Lingle, 2014). These approaches were complemented by patch-clamp experiments, and by measuring the cellular ion homeostasis and metabolism with fluorescence live-cell imaging, extracellular flux analysis and liquid chromatography-mass spectrometry (LC-MS)-based approaches.

Our results emphasize that the presence of BK_Ca_ in the IMM affects mitochondrial bioenergetics, thereby increasing BCC malignancy. This effect was specifically mediated by the DEC isoform of BK_Ca_, i.e., mitoBK_Ca_. Functional expression of BK_Ca_-DEC was validated by single-channel patch-clamp recordings BCC-derived mitoplasts. Importantly, we also identified BK_Ca_-DEC expression in a subset of BC patient biopsies using nanostring-based mRNA expression analysis. Finally, mitoBK_Ca_ crucially contributed to murine and human BCC proliferation and hypoxic resistance, and its activity increased lactate secretion resulting in higher extracellular acidification rates, even in the presence of O_2_.

Combined, our analyses provide, for the first time, a mechanistic link between functionally relevant mitochondrial BK_Ca_ isoforms in cancer cells and the promotion of the Warburg effect.

## Results

### Functional characterization of BK_Ca_ expression in BCCs

First, we assessed the functional expression of BK_Ca_ in established human BCC lines by performing whole-cell patch-clamp experiments. Primary BCCs obtained from transgenic, BC-bearing MMTV-PyMT WT or BK-KO mice were used as positive or negative controls, respectively (Mohr et al., 2022). Patch-clamp experiments revealed that depolarizing stimuli delivered in 20 mV increments induced K^+^ outward currents that were larger in MMTV-PyMT WT compared to BK-KO cells (**Figs. 1A** and **1B**, and **Figs. S1A** and **S1B**). To validate that these increased currents were elicited by BK_Ca_, we pharmacologically inhibited the channel by using either paxilline or iberiotoxin (Candia et al., 1992; Zhou and Lingle, 2014). While the current remained unaffected by these treatments in MMTV-PyMT BK-KO cells (**Fig. 1B** and **Fig. S1B**), peak currents (I_max_) were drastically reduced in MMTV-PyMT WT cells (**Fig. 1A** and **Fig. S1A**). Further, we analyzed the plasma membrane potential (ΔΨ_PM_) in these cells using the voltage-sensitive fluorescent dye Dibac4(3) (Adams and Levin, 2012) as well as current clamp experiments. The ΔΨ_PM_ was more polarized in MMTV-PyMT WT compared to BK-KO cells (**Fig. 1C and Fig. S1C**), as expected, due to the presence of hyperpolarizing BK_Ca_ channels in WT cells (N’Gouemo, 2014). Current clamp experiments unveiled a basal ΔΨ_PM_ of -44.0 ± 2.4 mV and -32.2 ± 2.1 mV for MMTV-PyMT WT and BK-KO cells, respectively (**Fig. S1C**). Iberiotoxin treatment equalized the ΔΨ_PM_ of the two genotypes (**Fig. S1C**). In line with these findings, using a recently developed, genetically encoded K^+^ sensor, NES lc-LysM GEPII 1.0 (Bischof et al., 2017), we found reduced cytosolic K^+^ concentrations ([K^+^]_cyto_) in MMTV-PyMT WT cells under basal conditions (83.07 ± 5.8 mM for WT ctrl vs. 121.2 ± 7.1 mM for BK-KO ctrl), which increased to the BK-KO cell level in response to iberiotoxin treatment (109.8 ± 10.8 mM for WT + IBTX vs. 114.6 ± 9.8 mM for BK-KO + IBTX) (**Figs. S1D** and **Fig. S1E**), explaining the PM depolarization. Subsequently, we recorded current-voltage relationships using the human BCC lines MDA-MB-453 and MCF-7, expressing either high or low levels of BK_Ca_ mRNA transcripts, respectively (Mohr et al., 2022). Analysis of the outward currents activated by depolarization demonstrated a paxilline- and iberiotoxin-sensitive current in MDA-MB-453 cells (**Fig. 1D** and **Fig. S1F**), indicative for functional BK_Ca_ channels in their PM, but not in MCF-7 cells (**Fig. 1E** and **Fig. S1G**), which is well in line with previous studies (Mohr et al., 2022). Based on the almost non-existent level of BK_Ca_-mediated PM currents in MCF-7 cells, we utilized these cells to express different BK_Ca_ isoforms fused to a red fluorescent protein (RFP). Therefore, we either used an earlier identified mitoBK_Ca_ isoform (Singh et al., 2013), namely BK_Ca_-DEC^RFP^, or the same channel lacking C-terminal amino acids including the DEC exon, hereinafter referred to as BK_Ca_^RFP^ (**Fig. 1F**). We hypothesized, that, in analogy to cardiac myocytes (Singh et al., 2013), the expression of BK_Ca_-DEC^RFP^ in MCF-7 cells may yield a functional channel present in the IMM. To test this, MCF-7 cells were first transiently transfected either with RFP fused to a glycosylphosphatidylinositol (GPI)-anchor (RFP-GPI) directing RFP to the PM, or linked to a cytochrome c oxidase subunit 8 (COX8) mitochondrial leading sequence, yielding a mitochondrial targeted fusion protein (mtRFP) (**Fig. S1H** and grey dotted lines in **Fig. 1G**). Analysis of the colocalization of mtRFP with MitoGREEN, a dye that specifically stains the mitochondrial matrix, resulted in high colocalization scores, while a low overlap was observed for RFP-GPI transfected MCF-7 cells (**Fig. S1H** and **Fig. 1G**, grey dotted lines). Subsequently, the same experiments were performed with MCF-7 cells expressing BK_Ca_^RFP^ or BK_Ca_-DEC^RFP^. Although the mitochondrial localization score was lower for BK_Ca_-DEC^RFP^ compared to mtRFP (compare **Fig. 1G** and **Fig. S1H**), this BK_Ca_ variant showed a significantly higher overlap with the mitochondrial dye than BK_Ca_^RFP^ (**Fig. 1G and Fig. S1I**). Despite rather moderate colocalization scores of BK_Ca_-DEC^RFP^ with MitoGREEN, on the level of single mitochondria, the RFP signal derived from BK_Ca_-DEC^RFP^ surrounded the MitoGREEN fluorescence signal originating from the mitochondrial matrix, as expected for a K^+^ channel located in the IMM (**Fig.1G** and **Fig. S1I**). As not all RFP signal in the BK_Ca_-DEC^RFP^ overexpressing MCF-7 originated from mitochondria, we investigated the PM localization of the two channel isoforms by patch-clamp. Compared to native MCF-7 cells (**Fig. 1E**), both, BK_Ca_^RFP^ and BK_Ca_-DEC^RFP^, increased the PM outward current (**Fig. 1H** and **Figs. S1J** and S**1K**). Despite comparable expression levels of the RFP signals (**Fig. 1L**), presence of BK_Ca_^RFP^ caused significantly bigger currents across the PM compared to BK_Ca_-DEC^RFP^ (**Fig. 1H**), indicative either for i) higher intracellular abundance or ii) major functional differences of BK_Ca_-DEC^RFP^. Importantly, the conductance of both channels was sensitive to the BK_Ca_ blockers paxilline and iberiotoxin, as treatment of the cells with both compounds resulted in PM conductance values comparable to those of native MCF-7 cells (**Figs. 1E** and **1H** and **Figs. S1J** and **S1K**).

**Figure 1:**
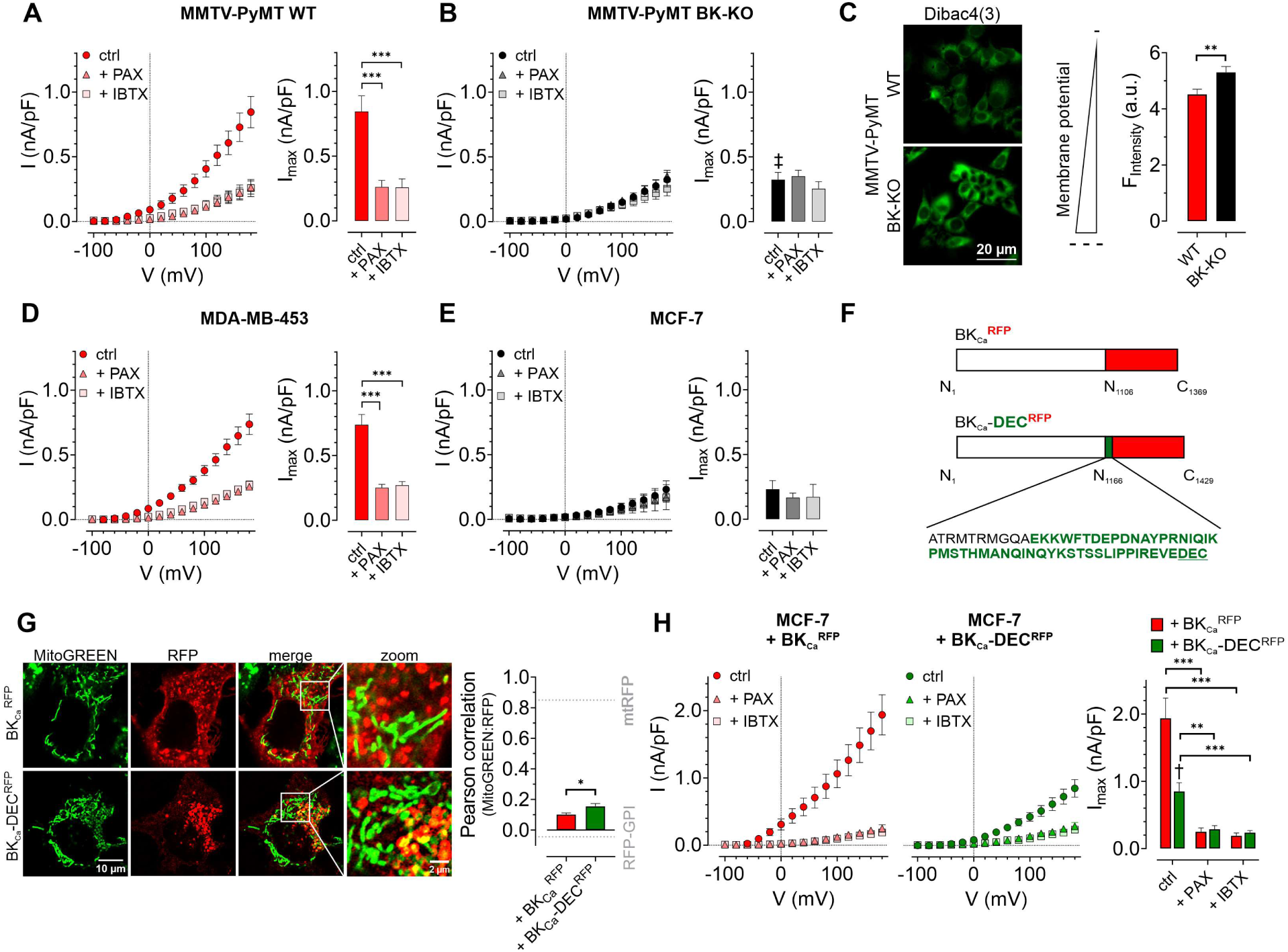
Characterization of BKCa channels in murine and human BCCs. (**A; B**) I-V curves (left) and corresponding maximal currents (right) of MMTV-PyMT WT (**A**) and MMTV-PyMT BK-KO cells (**B**), either under control conditions, or in the presence of paxilline or iberiotoxin. Data represents average ± SEM. n (cells) = 15 WT ctrl, 17 WT + PAX, 17 WT + IBTX, 16 BK-KO ctrl, 17 BK-KO + PAX, 19 BK-KO + IBTX. ***p≤0.001, Brown-Forsythe and Welch ANOVA test followed by Games-Howell’s multiple comparison test. ‡p≤0.001 compared to respective WT condition, Welch’s t-test. (**C**) Representative fluorescence images (left) and statistics (right) of MMTV-PyMT WT and BK-KO cells loaded with the ΔΨPM sensitive dye Dibac4(3). N = 6 independent experiments, **p≤0.01, Unpaired t-test. (**D**) I-V curves (left) and maximal currents (right) of MDA-MB-453 cells, either under control conditions, or in the presence of paxilline or iberiotoxin. Data represents average ± SEM. n (cells) = 30 ctrl, 22 + PAX, 24 + IBTX. ***p≤0.001, Kruskal-Wallis test followed by Dunn’s multiple comparison test. (**E**) I-V curves (left) and maximal currents (right) of MCF-7 cells, either under control conditions, or in the presence of paxilline or iberiotoxin. Data shows average ± SEM. n (cells) = 16 ctrl, 20 + PAX, 15 + IBTX. (**F**) Schematic representation of constructs used for over-expression in MCF-7 cells. The DEC exon is indicated in green. (**G**) Representative images (left) of MCF-7 cells either expressing BKCa^RFP^ (upper) or BKCa-DEC^RFP^ (lower), additionally stained with MitoGREEN. Average Pearson correlations ± SEM of MitoGREEN and RFP of BKCa or BKCa-DEC are shown. n (cells) = 17 BKCa – RFP, 22 BKCa-DEC^RFP^. *p≤0.05, Unpaired t-test. (**H**) I-V curves (left and middle) and corresponding maximal currents (right) of MCF-7 cells expressing BKCa (left) or BKCa-DEC (middle), respectively, either under control conditions, or in the presence of paxilline or iberiotoxin. Data represents average ± SEM. n (cells) = 18 BKCa^RFP^ ctrl, 14 BKCa^RFP^ + PAX, 19 BKCa^RFP^ + IBTX, 18 BKCa-DEC^RFP^ ctrl, 21 BKCa-DEC^RFP^ + PAX, 18 BKCa-DEC^RFP^ + IBTX. **p≤0.01, ***p≤0.001, Brown-Forsythe and Welch ANOVA test followed by Games-Howell’s multiple comparison test. †p≤0.01 between ctrl conditions, Welch’s t-test.

### BK_Ca_ modulates global and subcellular Ca^2+^ homeostasis in BCCs

As BK_Ca_ potentially affects cellular Ca^2+^ fluxes (Ouadid-Ahidouch and Ahidouch, 2013), we next investigated the cytosolic (**Figs. 2A-C**), endoplasmic reticulum (ER) (**Figs. 2D-F**), and mitochondrial (**Figs. 2G-I**) Ca^2+^ homeostasis in these cells. First, we assessed changes in the cytosolic Ca^2+^ concentration ([Ca^2+^]_cyto_) over-time in response to cell stimulation with the purinergic receptor agonist adenosine-5’-triphosphate (ATP) (Müller et al., 2020) using the fluorescent Ca^2+^ indicator Fura-2 (Grynkiewicz et al., 1985). Of note, extracellular ATP, released for example from necrotic cells in the tumor microenvironment, could be of pathophysiological relevance for BCCs (Di Virgilio et al., 2018), and all cell lines used in our study were previously reported to respond to ATP stimulation with a significant increase in intracellular Ca^2+^ (Chadet et al., 2014; Gross et al., 2022; Klijn et al., 2015). These experiments were either performed under control conditions (**Fig. 2A**) or in the presence of paxilline (**Fig. S2A**) or iberiotoxin (**Fig. S2B**). Analysis of the Fura-2 fluorescence emission ratio showed significantly elevated basal (**Fig. 2A**) and ATP-elicited maximal [Ca^2+^]_cyto_ responses (**Fig. S2C**) in MMTV-PyMT WT compared to BK-KO cells under control conditions. Interestingly, the elevated basal [Ca^2+^]_cyto_ of WT cells was reduced to the BK-KO cell level in response to paxilline (**Fig. 2A** and **Fig. S2A**), but not by iberiotoxin (**Fig. 2A** and **Fig. S2B**), which is a peptide-based pore-blocking toxin that cannot pass the PM (Candia et al., 1992). To validate the observed basal differences in [Ca^2+^]_cyto_, we additionally performed ionomycin-based calibrations by chelating intracellular Ca^2+^, followed by Fura-2 saturation (**Fig. S2F-H**). These experiments confirmed the observed differences in basal [Ca^2+^]_cyto_, with [Ca^2+^]_cyto_ of 170.2 ± 8.0 nM, 98.40 ± 4.8 nM and 178.5 ± 13.5 nM for MMTV-PyMT WT, and 102.3 ± 4.9 nM, 96.49 ± 3.3 nM and 114.9 ± 4.3 nM for MMTV-PyMT BK-KO cells, under control conditions, or in the presence of paxilline or iberiotoxin, respectively. Subsequently, basal and ATP-elicited maximal [Ca^2+^]_cyto_ transients were recorded in the human BCC line MDA-MB-453, where both BK_Ca_ blockers reduced the basal [Ca^2+^]_cyto_ (**Fig. 2B**), while maximal [Ca^2+^]_cyto_ was not altered by these treatments (**Fig. S2D**). Again, ionomycin-based experiments confirmed the BK_Ca_-modulation dependent reduction of [Ca^2+^]_cyto_ (**Fig. S2I**), with [Ca^2+^]_cyto_ of 110.7 ± 8.0 nM, 64.1 ± 4.8 nM and 78.2 ± 13.5 nM under control, paxilline- or iberiotoxin-treated conditions, respectively. Eventually, [Ca^2+^]_cyto_ was assessed in MCF-7 cells expressing or lacking the different BK_Ca_ isoforms (**Fig. 2C**). These experiments confirmed previous findings with the other BCC lines, as the expression of both splice variants increased the basal (**Fig. 2C** and **Fig. S2J**) and maximal [Ca^2+^]_cyto_ (**Fig. S2E**). [Ca^2+^]_cyto_ in MCF-7 cells was found to be 31.5 ± 3.7 nM, 58.1 ± 6.4 nM and 52,9 ± 5.6 nM in MCF-7 control cells expressing RFP, or MCF-7 cells expressing BK_Ca_^RFP^ or BK_Ca_-DEC^RFP^, respectively.

**Figure 2:**
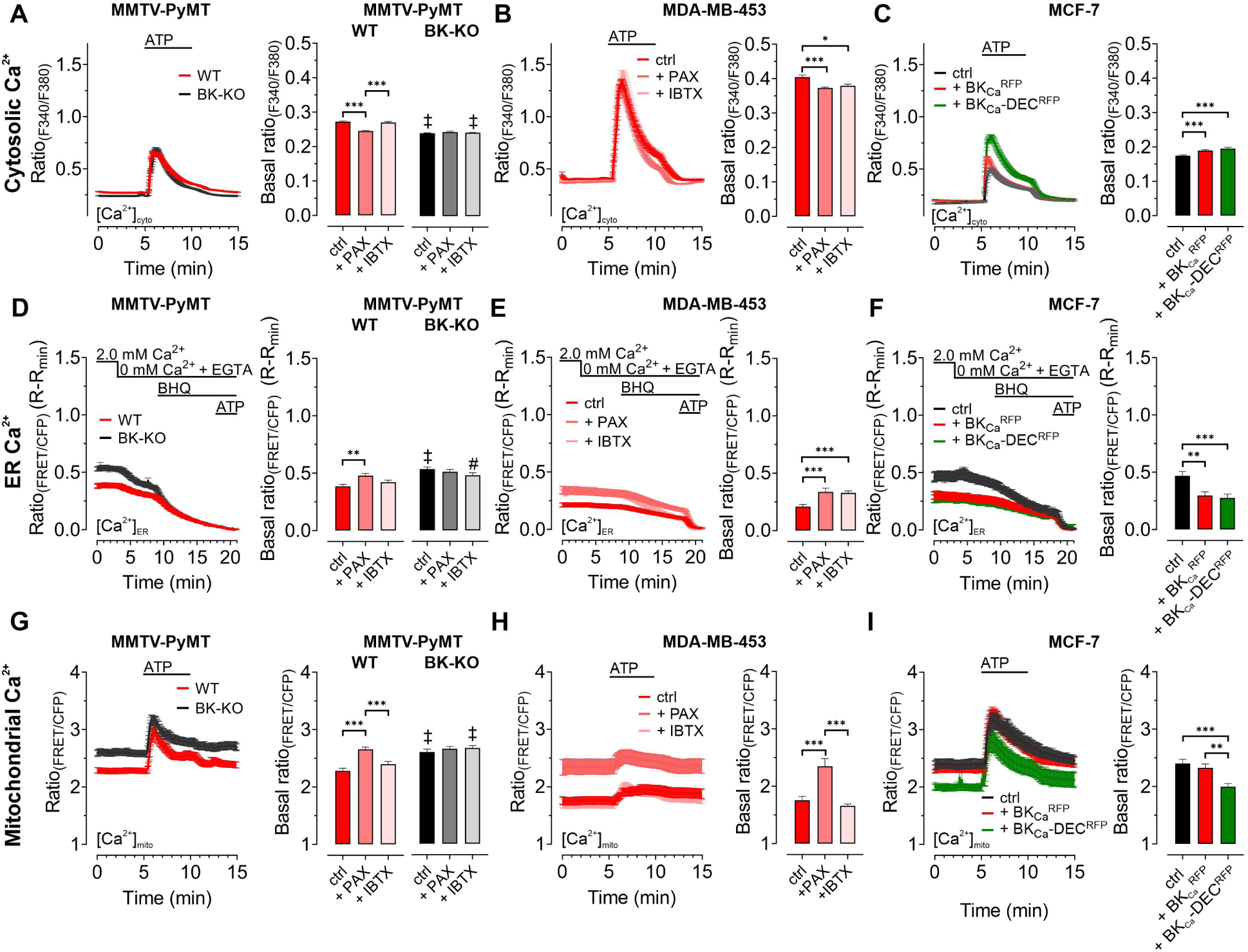
BKCa modulates the subcellular Ca^2+^ homeostasis in BCCs. Cytosolic (**A – C**), endoplasmic reticulum (ER) (**D – F**), and mitochondrial Ca^2+^ dynamics (**G – I**) over-time of MMTV-PyMT WT and MMTV-PyMT BK-KO cells (**A, D, G**), MDA-MB-453 cells (**B, E, H**) or MCF-7 cells (**C, F, I**). All data represent average ± SEM. At time points indicated in the panels, cytosolic and mitochondrial Ca^2+^ alterations were evoked by extracellular stimulation with ATP (**A – C, G – I**), or by Ca^2+^ depletion of the ER using Ca^2+^-free buffer containing the Ca^2+^ chelator EGTA, followed by administration of the SERCA inhibitor BHQ prior to ATP administration (**D – F**). MMTV-PyMT (**A, D, G**) and MDA-MB-453 (**B, E, H**) cells were either measured under control conditions, or in the presence of paxilline (+ PAX) or iberiotoxin (+ IBTX). MCF-7 cells (**C, F, I**) either expressed an RFP (ctrl), BKCa^RFP^, or BKCa-DEC^RFP^. N (independent experiments) / n (cells analyzed) = **A:** 17/784 WT ctrl, 18/857 BK-KO ctrl, 6/300 WT + PAX, 6/300 BK-KO + PAX, 5/318 WT + IBTX, 5/304 BK-KO + IBTX, **B:** 4/151 ctrl, 4/132 + PAX, 4/87 + IBTX, **C:** 5/111 ctrl, 5/117 + BKCa^RFP^, 5/91 + BKCa-DEC^RFP^, **D:** 14/116 WT ctrl, 13/117 BK-KO ctrl, 8/71 WT + PAX, 8/92 BK-KO + PAX, 6/102 WT + IBTX, 6/86 BK-KO + IBTX, **E:** 7/44 ctrl, 9/34 + PAX, 5/49 + IBTX, **F:** 4/25 ctrl, 4/35 + BKC^aRFP^, 4/38 + BKCa-DEC^RFP^, **G:** 11/47 WT ctrl, 12/86 BK-KO ctrl, 6/46 WT + PAX, 6/58 BK-KO + PAX, 5/59 WT + IBTX, 4/43 BK-KO + IBTX, **H:** 8/33 ctrl, 8/28 + PAX, 5/22 + IBTX, **I:** 5/28 ctrl, 4/27 + BKCa^RFP^, 4/24 + BKCa-DEC^RFP^. **p≤0.01, ***p≤0.001. Kruskal-Wallis test followed by Dunn’s MC test (**A, B, C, D, I**), One-Way ANOVA test followed by Tukey’s MC test (**E, F, G**) or Brown-Forsythe ANOVA test followed by Games-Howell’s MC test (**H**). #p≤0.05, ‡p≤0.001 compared to respective WT condition, Mann-Whitney test (**A, D,** ctrl in **G**), or Welch’s t test (+ IBTX in **G**).

Next, we visualized changes in the ER [Ca^2+^] ([Ca^2+^]_ER_). Therefore, BCCs were transfected with D1ER, an established genetically encoded, FRET-based Ca^2+^ sensor targeted to the lumen of the ER (Palmer et al., 2004). [Ca^2+^]_ER_ was depleted by extracellular Ca^2+^ removal and chelation by EGTA, followed by inhibition of the sarcoplasmic endoplasmic reticulum Ca^2+^ ATPase (SERCA) with BHQ and activation of inositol 1,4,5-trisphosphate (IP_3_) receptors upon purinergic receptor stimulation with ATP (Lape et al., 2008; Müller et al., 2020; Salter and Hicks, 1995). Experiments were either performed under control conditions (**Figs. 2D-F**), or in the presence of paxilline or iberiotoxin for MMTV-PyMT (**Fig. 2D** and **Figs. S2K** and **S2L),** MDA-MB-453 (**Fig. 2E**), and MCF-7 cells expressing BK_Ca_^RFP^ or BK_Ca_-DEC^RFP^ (**Fig. 2F**). Throughout all BCCs investigated, expression of BK_Ca_ reduced [Ca^2+^]_ER_, potentially indicating that the channel was i.) functional in this cell compartment and ii.) involved in regulating the Ca^2+^ homeostasis of the ER. In MMTV-PyMT WT cells, [Ca^2+^]_ER_ was restored by the cell-permeable BK_Ca_ inhibitor paxilline (**Fig. 2D** and **Fig. S2K**) but not by the cell-impermeable iberiotoxin (**Fig. 2D** and **Fig. S2L**), while both inhibitors restored the [Ca^2+^]_ER_ in MDA-MB-453 cells (**Fig. 2E**). Accordingly, overexpression of both RFP-tagged BK_Ca_ isoforms in MCF-7 cells depleted the [Ca^2+^]_ER_ (**Fig. 2F**).

[Ca^2+^]_cyto_ and [Ca^2+^]_ER_ reportedly affect the mitochondrial [Ca^2+^] ([Ca^2+^]_mito_) (Wacquier et al., 2019). Since an accumulation of K^+^ within the mitochondrial matrix may oppose mitochondrial Ca^2+^ uptake (Checchetto et al., 2021), we assessed the effects of functional BK_Ca_ expression on [Ca^2+^]_mito_ utilizing 4mtD3cpV, a genetically encoded Ca^2+^ sensor targeted to the mitochondrial matrix (Palmer et al., 2006). Again, BCCs were treated with ATP to investigate the [Ca^2+^]_mito_ upon cell stimulation. In MMTV-PyMT WT (**Fig. 2G**) as well as MDA-MB-453 cells (**Fig. 2H**), functional expression of BK_Ca_ reduced basal [Ca^2+^]_mito_. Interestingly, in these two BCC types, basal and maximally elicited [Ca^2+^]_mito_ peaks increased in response to paxilline (**Figs. 2G** and **2H** and **Figs. S2M-P**), but not iberiotoxin treatment (**Figs. 2G** and **2H** and **Figs. S2M-P**). These findings confirm a role of intracellular located BK_Ca_ channels in modulating [Ca^2+^]_mito_ dynamics. Importantly, in MCF-7 cells, basal [Ca^2+^]_mito_ was only affected upon expression of the mitochondrially targeted BK_Ca_-DEC^RFP^, but not BK_Ca_ ^RFP^ (**Fig. 2I**), whereas neither of the two isoforms affected the maximal [Ca^2+^]_mito_ (**Fig. S2Q**). Presumably, by facilitating K^+^ fluxes across the IMM, BK_Ca_-DEC^RFP^ expression reduces the driving force for Ca^2+^ uptake and thus the resulting Ca^2+^ signals in the mitochondrial matrix. Basal differences in all of the utilized cell lines under the different conditions were additionally validated by ionomycin treatment, followed by intracellular Ca^2+^ chelation (**Figs. S2R-V**).

In total, our [Ca^2+^] imaging approaches emphasize that different BK_Ca_ isoforms at different localizations, i.e. the PM or intracellular organelles, may either amplify or weaken [Ca^2+^]_cyto_ and [Ca^2+^]_ER_ signals, respectively. These opposing effects are expected, as BK_Ca_-mediated uptake of K^+^ into the ER should limit the Ca^2+^ uptake capacity of this subcellular compartment, while functional channel expression at the PM might increase the driving force for Ca^2+^ influx to fuel [Ca^2+^]_cyto_. [Ca^2+^]_mito_, however, seems to be exclusively and effectively suppressed by the activity of intracellularly located BK_Ca_ variants in MMTV-PyMT WT and MDA-MB-453 cells and solely by the expression of BK_Ca_-DEC^RFP^ in MCF-7 cells.

### The metabolic activity of murine and human BCCs is modulated by intracellular BK_Ca_

Given the involvement of Ca^2+^ and K^+^ in regulating key features of cellular metabolism (Bischof et al., 2021; Dejos et al., 2020; Gohara and Di Cera, 2016), we proposed, that BK_Ca_ may alter metabolic activities of BCCs (Rossi et al., 2019). Hence, we analyzed the extracellular acidification rate (ECAR) as a measure of lactate secretion, i.e. glycolytic activity in MMTV-PyMT and MDA-MB-453 cells (Burgstaller et al., 2021a). ECAR measurements unveiled an increased basal ECAR of MMTV-PyMT WT compared to BK-KO cells (**Figs. 3A** and **3B**). Subsequently, we performed a mitochondrial stress test by injection of Oligomycin-A, FCCP, and Antimycin-A, which increased ECARs in both cell types due to ATP-synthase inhibition, mitochondrial uncoupling and complex III blockade, respectively (**Fig. 3A**). Maximal ECAR, received upon FCCP injection, was elevated in WT compared to BK-KO cells (**Fig. 3C**). Besides this evidence derived from a gene-targeted BK_Ca_ channel-deficient model, we used paxilline or iberiotoxin as pharmacological BK_Ca_ modulators (**Figs. S3A** and **S3B**). Interestingly, only paxilline, but not iberiotoxin treatment equalized basal (**Fig. 3B**) and maximal ECARs (**Fig. 3C**) between WT and BK-KO, suggesting that intracellular BK_Ca_ channels stimulate the glycolytic activity of BCCs. Moreover, in MDA-MB-453 cells iberiotoxin treatment did neither affect basal (**Fig. 3D** and **3E**) nor maximal ECARs (**Fig. 3F**). Contrary to MMTV-PyMT WT cells, however, paxilline increased basal and maximal ECAR in these cells (**Figs. 3E** and **3F**), suggesting that the cell lines examined strongly differ in their metabolic settings or in other H^+^ releasing processes contributing to extracellular acidification.

**Figure 3:**
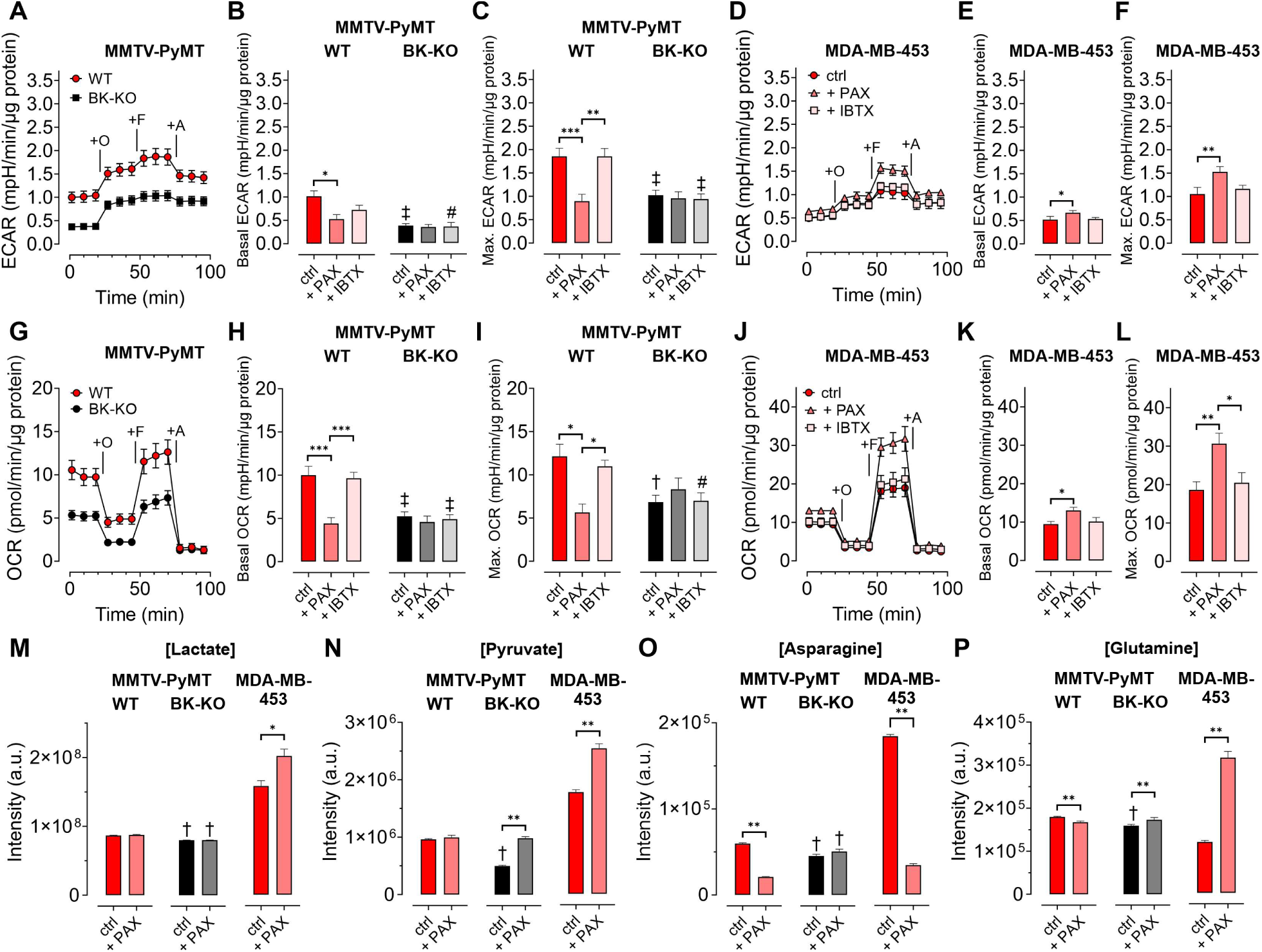
BKCa channels alter the metabolic activity of BCCs. (**A, D**) Average ECAR over-time ± SEM of MMTV-PyMT WT (**A,** red) and BK-KO cells (**A,** black), or MDA-MB-453 cells (**D**) in response to administration of Oligomycin-A (+O), FCCP (+F) and Antimycin-A (+A) at time points indicated. (**B, E**) Average basal and (**C, F**) maximal ECAR ± SEM of MMTV-PyMT WT (**B, C,** left) and BK-KO cells (**B, C,** right), or MDA-MB-453 cells (**E, F**) under control conditions, or in the presence of paxilline or iberiotoxin. (**G, J**) Average OCR over-time ± SEM of MMTV-PyMT WT (**G,** red) and BK-KO cells (**G,** black), or MDA-MB-453 cells (**J**) in response to administration of Oligomycin-A (+O), FCCP (+F) and Antimycin-A (+A) at time points indicated. (**H, K**) Average basal and (**I, L**) maximal OCR ± SEM of MMTV-PyMT WT (**H, I,** left) and BK-KO cells (**H, I,** right), or MDA-MB-453 cells (**K, L**) under control conditions, or in the presence of paxilline or iberiotoxin. (**M – P**) LC-MS-based determination of major glycolytic and mitochondrial metabolites in particular Lactate (**M**, [Lactate]), Pyruvate (**N**, [Pyruvate]), Asparagine (**O**, [Asparagine]) and Glutamine (**P**, [Glutamine]) of MMTV-PyMT WT (left panels), BK-KO (middle panels) and MDA-MB-453 cells (right panels), either under control conditions or after cell cultivation with paxilline. N (independent experiments) = **A, B, C, G, H, I:** 7 WT ctrl and BK-KO, 3 for all others, **D, E, F, J, K, L:** 3 for all, **M – P:** 7 for BK-KO ctrl, 6 for all others. *p≤0.05, **p≤0.01, ***p≤0.001, Kruskal-Wallis test followed by Dunn’s MC test (**B, E, F, I**), Brown-Forsythe and Welch ANOVA test followed by Games-Howell’s MC test (**C, H**), One-Way ANOVA test followed by Tukey’s MC test (**K, L**) or Mann-Whitney test (**M – P**). #p≤0.05, †p≤0.01, ‡p≤0.001, to respective WT condition, Mann-Whitney test (**B, C,** + IBTX in **I**, **M – P**), Welch’s t-test (ctrl in **H** and **I**) or Unpaired t-test (+ IBTX in **H**).

Next, we investigated the oxygen consumption rates (OCRs) (Burgstaller et al., 2021a). (**Fig. 3G**). Interestingly, the increased basal and maximal ECAR (**Figs. 3A-C**) in MMTV-PyMT WT cells correlated with a higher basal (**Fig. 3H**) and maximal (**Fig. 3I**) OCR compared to BCCs lacking BK_Ca_. Paxilline treatment reduced the basal and maximal OCR of BK_Ca_ proficient MMTV-PyMT cells to the BK-KO level (**Figs. 3H** and **3I** and **Fig. S3C**), while iberiotoxin did not have any effect (**Figs. 3H** and **3I** and **Fig. S3D**) again suggesting that that the latter toxin cannot reach the metabolism-relevant population of intracellular BK_Ca_ channels. Again, in MDA-MB-453 cells, paxilline treatment had an opposite effect to the observations that were made in MMTV-PyMT WT cells, as paxilline treatment increased the OCR (**Figs. 3J-L**), potentially due to the increased supply of oxidative phosphorylation with glycolytic substrates (**Figs. 3D-F**). More importantly, regardless of this difference between the BCC lines iberiotoxin treatment did consistently not affect the OCR of MDA-MB-453 cells (**Figs. 3J-L**).

To confirm that BK_Ca_ regulates the bioenergetic profile of BCCs, we subsequently applied LC-MS-based metabolomics according to the workflow presented in **Figure S3E**. We included typical analytes of glycolysis such as lactate (**Fig. 3M**) and pyruvate (**Fig. 3N**), as well as selected metabolites of mitochondrial metabolism including asparagine (**Fig. 3O**) and glutamine (**Fig. 3P**). Single time-point measurements confirmed the metabolic differences between MMTV-PyMT WT and BK-KO cells and further demonstrated that BK_Ca_ inhibition by paxilline directly affected the concentrations of these metabolites (**Figs. 3M-P**). As observed before by the ECAR and OCR measurements, MMTV-PyMT and MDA-MB-453 cells responded, however, differently to paxilline treatment (**Figs. 3M-P**). In summary, our data strengthen the notion that intracellular BK_Ca_ modulates the cellular energy balance of murine and human BCCs.

### BK_Ca_ alters the mitochondrial function of BCCs

To clarify how BK_Ca_ regulates BCC cell metabolic activities, we examined cellular bioenergetics in real time using single-cell fluorescence microscopy. First, we assessed the mitochondrial membrane potential (ΔΨ_mito_) of MMTV-PyMT and MDA-MB-453 cells using TMRM under control and BK_Ca_ inhibitor-treated conditions, followed by depolarization of ΔΨ_mito_ using the proton ionophore FCCP (Joshi and Bakowska, 2011). These measurements unveiled, however, a less polarized ΔΨ_mito_ of MMTV-PyMT WT cells compared to BK-KO cells under control conditions (**Figs. 4A** and **4B**). Paxilline, but not iberiotoxin, equalized ΔΨ_mito_ between MMTV-PyMT WT and BK-KO cells (**Fig. 4B**). Identical results were obtained in MDA-MB-453 cells (**Figs. 4C** and **4D**).

**Figure 4:**
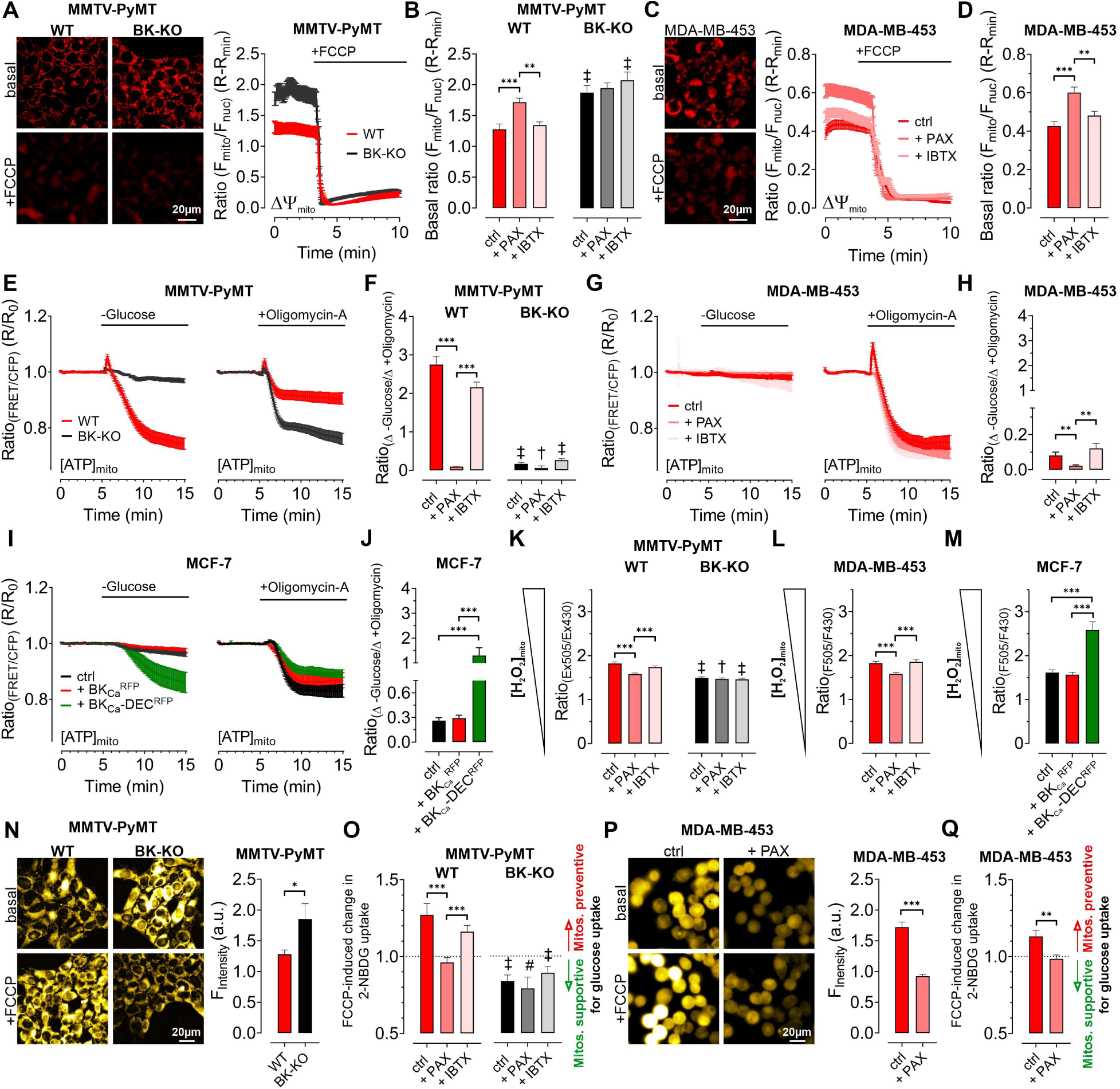
Expression of BKCa modulates mitochondrial function and glucose uptake of BCCs. (**A – D**) Representative fluorescence images and -ratios (Fmito/Fnuc) over-time (**A, C**), and corresponding statistics ± SEM (**B, D**) representing ΔΨmito of TMRM-loaded MMTV-PyMT WT and BK-KO (**A, B**) and MDA-MB-453 cells (**C, D**) under basal conditions (**A, C**, upper images) and upon administration of FCCP for mitochondrial depolarization (**A, C** lower images). (**E – J**) [ATP]mito dynamics ± SEM over-time of MMTV-PyMT WT and BK-KO cells (**E**), MDA-MB-453 cells (**G**) and MCF-7 cells (**I**) in response to extracellular glucose removal (left panels) or upon administration of Oligomycin-A (right panels). **F, H** and **J** show changes of [ATP]mito induced by glucose removal to Oligomycin-A administration ± SEM, under control conditions, or in the presence of paxilline or iberiotoxin (**F, H**), or upon expression of BKCa^RFP^ or BKCa-DEC^RFP^ (**J**). (**K – M**) Basal mitochondrial H2O2 concentrations ± SEM of MMTV-PyMT WT (**K**, left), BK-KO (**K**, right), MDA-MB-453 (**L**) and MCF-7 cells (**M**), either under control conditions, in the presence of paxilline or iberiotoxin (**K, L**), or upon expression of BKCa^RFP^ or BKCa-DEC^RFP^ (**M**). (**N, P**) Representative fluorescence wide-field images (left) and corresponding statistics ± SEM (right) of MMTV-PyMT WT (**N,** left images and red bars) and BK-KO cells (**N,** right images and black bars) or MDA-MB-453 cells (**P**) incubated with 2-NBDG, either in the absence (upper images) or presence of FCCP (lower images). (**O, Q**) Average ± SEM of FCCP induced change in 2-NBDG uptake of MMTV-PyMT WT (**O**, left) and BK-KO cells (**O**, right), or MDA-MB-453 cells (**Q**) either under control conditions, or in the presence of paxilline or iberiotoxin. Values above 1 indicate that mitochondria prevent, values below 1 that mitochondria support glucose uptake. N (independent experiments) / n (cells analyzed) = **A**, **B:** 4/75 WT ctrl, 4/90 WT + PAX, 4/86 WT + IBTX, 4/91 BK-KO ctrl, 4/89 BK-KO + PAX, 4/100 BK-KO + IBTX, **C, D:** 4/113 ctrl, 4/97 + PAX, 4/103 + IBTX, **E, F:** [-Glucose]: 8/55 WT ctrl, 6/45 WT + PAX, 7/27 WT + IBTX, 8/65 BK-KO ctrl, 6/57 BK-KO + PAX, 7/28 BK-KO + IBTX, [+ Oligomycin-A]: 11/52 WT ctrl, 7/53 WT + PAX, 7/34 WT + IBTX, 8/87 BK-KO ctrl, 6/35 BK-KO + PAX, 5/45 BK-KO + IBTX. **G, H:** [-Glucose]: 5/14 ctrl, 3/13 + PAX, 5/13 + IBTX, [+ Oligomycin-A]: 5/33 ctrl, 3/21 + PAX, 8/27 + IBTX, **I, J:** [-Glucose]: 6/48 ctrl, 5/23 + BK ^RFP^, 5/20 + BK -DEC^RFP^, [+ Oligomycin-A]: 5/27 ctrl, 5/23 + BK ^RFP^, 5/37 + BK -DEC^RFP^, **K:** 3/33 WT ctrl, 4/51 WT + PAX, 4/54 WT + IBTX, 4/55 BK-KO ctrl, 4/51 BK-KO + PAX, 4/54 BK-KO + IBTX, **L:** 4/31 ctrl, 4/39 + PAX, 4/31 + IBTX, **M:** 4/29 ctrl, 4/17 + BKCa^RFP^, 4/21 + BKCa-DEC^RFP^, **N – Q:** 4 for all. *p≤0.05, **p≤0.01, ***p≤0.001, Kruskal-Wallis test followed by Dunn’s MC test (**B, D, F, H, J, K, O**), Brown-Forsythe and Welch ANOVA test followed by Games-Howell’s MC test (**L, M**), Mann-Whitney test (**N**), Unpaired t-test (**P**) or Welch’s t-test (**Q**). #p≤0.05, †p≤0.01, ‡p≤0.001, to respective WT condition, Mann-Whitney test (**B, F,** + PAX and + IBTX in **K,** ctrl in **O**), Unpaired t-test (ctrl in **K,** + PAX and + IBTX in **O**).

Interestingly, we found that ΔΨ_mito_ of MMTV-PyMT cells was tightly dependent on the extracellular glucose concentration ([GLU]_ex_) (**Fig. S4A**). Under high [GLU]_ex_ conditions (25.0 mM), the difference in ΔΨ_mito_ between MMTV-PyMT WT and BK-KO cells disappeared, as ΔΨ_mito_ increased significantly in WT and decreased significantly in BK-KO cells compared to low [GLU]_ex_ (2.0 mM). Because F_O_F_1_-ATP-synthase (ATP Synthase) may hydrolyze ATP in an attempt to maintain ΔΨ_mito_, these results strongly suggest that the forward (ATP producing) *vs* reverse (ATP consuming) mode of the ATP synthase is affected by the BK_Ca_ status of the cells (Naguib et al., 2018).

To unravel the substrate dependency of these BCCs for maintaining their energy homeostasis in dependency of BK_Ca_, we measured the mitochondrial ATP concentration ([ATP]_mito_) using a genetically encoded ATP sensor targeted to the mitochondrial matrix, mtAT1.03 (Imamura et al., 2009). Mitochondrial ATP reportedly responds most dynamically to energy metabolism perturbations (Depaoli et al., 2018). To assess the sources of ATP in the cells, we either deprived the cells of [GLU]_ex_, or administered Oligomycin-A to inhibit the ATP-synthase (Depaoli et al., 2018) (**Figs. 4E-J** and **Figs. S4B-G**). In MMTV-PyMT cells, glucose removal as well as ATP-synthase inhibition reduced [ATP]_mito_, albeit the effect was more pronounced upon [GLU]_ex_ depletion (**Figs. 4E** and **4F** and **Figs. S4B** and **S4C**). Remarkably, paxilline, but not iberiotoxin treatment reduced the glucose, and increased ATP-synthase dependency of MMTV-PyMT WT cells for maintaining [ATP]_mito_ (**Figs. 4E** and **4F** and **Figs. S4B** and **S4C**). These observations were confirmed in MDA-MB-453 cells, even though i.) these cells showed an overall higher dependency on the ATP-synthase, and ii.) the paxilline-sensitivity of [ATP]_mito_ maintenance was much less pronounced (**Figs. 4G** and **4H** and **Figs. S4D** and **S4E**).

To validate these findings in a BK_Ca_ minimal to depleted model, these experiments were performed with MCF-7 cells either expressing exclusively RFP as control, BK_Ca_^RFP^, or BK_Ca_-DEC^RFP^ (**Fig. 4I**). Expression of BK_Ca-_DEC^RFP^, but not BK_Ca_^RFP^, resulted in a high dependency on [GLU]_ex_ to maintain [ATP]_mito_ while the [ATP]_mito_ rundown in the presence of Oligomycin A was identical for both BK_Ca_ splice variants. This suggests that BK_Ca_-DEC^RFP^ specifically triggers a high [GLU]_ex_ sensitivity and simultaneously an independency on ATP derived from ATP-synthase for maintaining [ATP]_mito_ as demonstrated by the ratio of the respective changes under these experimental conditions (**Figs. 4I** and **4J** and **Figs. S4F** and **S4G**). These findings suggest that intracellular (mitochondrial) BK_Ca_ contributes to the metabolic reprogramming of BCCs.

In an extension to extracellular flux analyses, single time-point LC-MS metabolomics (**Fig. 3**), and high-resolution live-cell imaging experiments (**Figs. 4A-J**), pointing to a BK_Ca_-dependent “oncometabolic” phenotype, we assessed the concentrations of mitochondrial hydrogen peroxide ([H_2_O_2_]_mito_) using mitoHyPer3 (Bilan et al., 2013), a genetically encoded fluorescent indicator for monitoring H_2_O_2_ in the mitochondrial matrix (**Figs. 4K-M**). These experiments demonstrated increased levels of [H_2_O_2_]_mito_ in MMTV-PyMT WT (**Fig. 4K**) and MDA-MB-453 cells (**Fig. 4L**), or specifically upon BK_Ca_-DEC^RFP^ expression in MCF-7 cells (**Fig. 4M**). Excessive reactive oxygen species (ROS) synthesis is caused by uncoupling of the respiratory chain and it is considered as an indicator of mitochondrial stress, which, among other reasons, promotes mutagenesis and BCC progression. In this regard, mitoBK_Ca_ may serve as an “uncoupling” protein (Gałecka et al., 2021), triggering ATP synthase to operate in reverse mode to consume instead of producing ATP, thereby counteracting the dissipation of the proton gradient and consequently the loss of ΔΨ_mito_. Consequently, the lack of ATP must be compensated, e.g. by accelerating glycolysis. To investigate this assumption, we performed glucose uptake measurements using 2-NBDG, a fluorescent glucose analogue, which is taken up via glucose transporters (GLUTs), phosphorylated by hexokinase (HK) isoforms to generate 2-NBDG-6-phosphate and subsequently remains within the cell (Bischof et al., 2021) (**Figs. S4H** and **S4I**). Unexpectedly, we found that MMTV-PyMT BK-KO cells showed a higher rate of glucose uptake and phosphorylation compared to WT cells under basal conditions (**Fig. 4N**). To unravel the role of mitochondria in contributing to glucose uptake by supplying ATP to HKs, we next performed these experiments in the presence of FCCP, which disrupts mitochondrial ATP production (Losano et al., 2017). If ATP-synthase works in forward mode, FCCP treatment should reduce or even prevent oxidative ATP production, and, subsequently, ATP-dependent 2-NBDG phosphorylation (**Fig. S4H**). Contrary, if ATP-synthase works in reverse mode, it may compete with HKs for ATP, and FCCP treatment should abolish this competition, leading to increased 2-NBDG phosphorylation in BCCs (**Fig. S4I**). Interestingly, our experiments unveiled increased 2-NBDG uptake in MMTV-PyMT WT and reduced uptake in BK-KO cells upon mitochondrial depolarization (**Fig. 4N** and **Figs. S4J** and **S4K**). To facilitate the interpretation, we calculated the FCCP-induced change in 2-NBDG uptake. These analyses revealed that mitochondria are less effective in assisting 2-NBDG uptake and phosphorylation in MMTV-PyMT WT cells, while they “support” these processes in BK-KO cells under control conditions (**Fig. 4O**). To test whether the changes in 2-NBDG uptake are sensitive to pharmacological modulation of BK_Ca_, we performed a similar set of experiments in the presence of paxilline or iberiotoxin. While iberiotoxin treatment did not have any effect, paxilline treatment shifted the activity of the ATP-synthase to the ATP-supplying “forward” mode (**Fig. 4O** and **Figs. S4J** and **S4K**). Comparable effects were obtained in MDA-MB-453, despite paxilline treatment reduced basal accumulation of 2-NBDG in these cells (**Figs. 4P** and **4Q** and **Fig. S4L**). Overall, the experiments performed so far suggest that mitochondria rather consume than generate ATP if BK_Ca_ was functionally expressed intracellularly in BCCs.

### BK_Ca_ locates functionally in the IMM of murine and human BCCs

So far, our data suggest an important contribution of intracellularly located BK_Ca_, possibly mitoBK_Ca_, in reprogramming cancer cell metabolism. Thus, we applied an electrophysiological approach to provide functional evidence for endogenous mitoBK_Ca_. Single-channel patch-clamp experiments were conducted using mitoplasts isolated from MMTV-PyMT WT and BK-KO, MDA-MB-453, and MCF-7 cells. In MDA-MB-453- and MMTV-PyMT WT-derived mitoplasts we indeed detected channels of large conductance (**Fig. 5A** and **Fig. S5A**). For these channels, the open probability did not significantly vary with the voltage (**Fig. 5B**). Only at very negative and positive voltages of approximately -150 mV and +150 mV differences in the open probabilities were observed (**Fig. 5C**). The bursts of single-channel openings showed an average conductance of 212 ± 2 pS (**Fig. 5D**). This large conductance and the sensitivity towards Ca^2+^-(**Figs. 5E** and **5F** and **Fig. S5B**) and paxilline (**Figs. 5E** and **5F** and **Fig. S5C**) pointed to mitoBK_Ca_ that exhibits the pharmacologic characteristics of canonical BK_Ca_ channels present at the PM. Overall, our electro-pharmacological experiments unveiled 3 different classes of channels with either small (≤100 pS), medium (≤150 pS) or large (∼210 pS) conductance, where the latter conductance corresponds to mitoBK_Ca_ (Liu et al., 1999). The lower conductance values may represent mitochondrial IK_Ca_, SK_Ca_, or K_ATP_ channel activity, but this was not investigated further in our study. Notably, mitoBK_Ca_ was detected at a frequency of 15% in 210 patches of MDA-MB-453 mitoplasts and at a lower frequency of 6% in 127 mitoplast patches of MMTV-PyMT WT cells. These numbers may, however, underestimate the actual abundance of the channel, as only a small area of the mitoplast is examined with each patch. Importantly, this channel was absent in 76 mitoplast patches from MMTV-PyMT BK-KO and 58 mitoplast patches from MCF-7 cells (**Fig. 5G**). These findings provide evidence that the molecular entity for the K^+^ channel derives from the nuclear *Kcnma1* gene, which is ablated in the MMTV-PyMT BK-KO.

**Figure 5:**
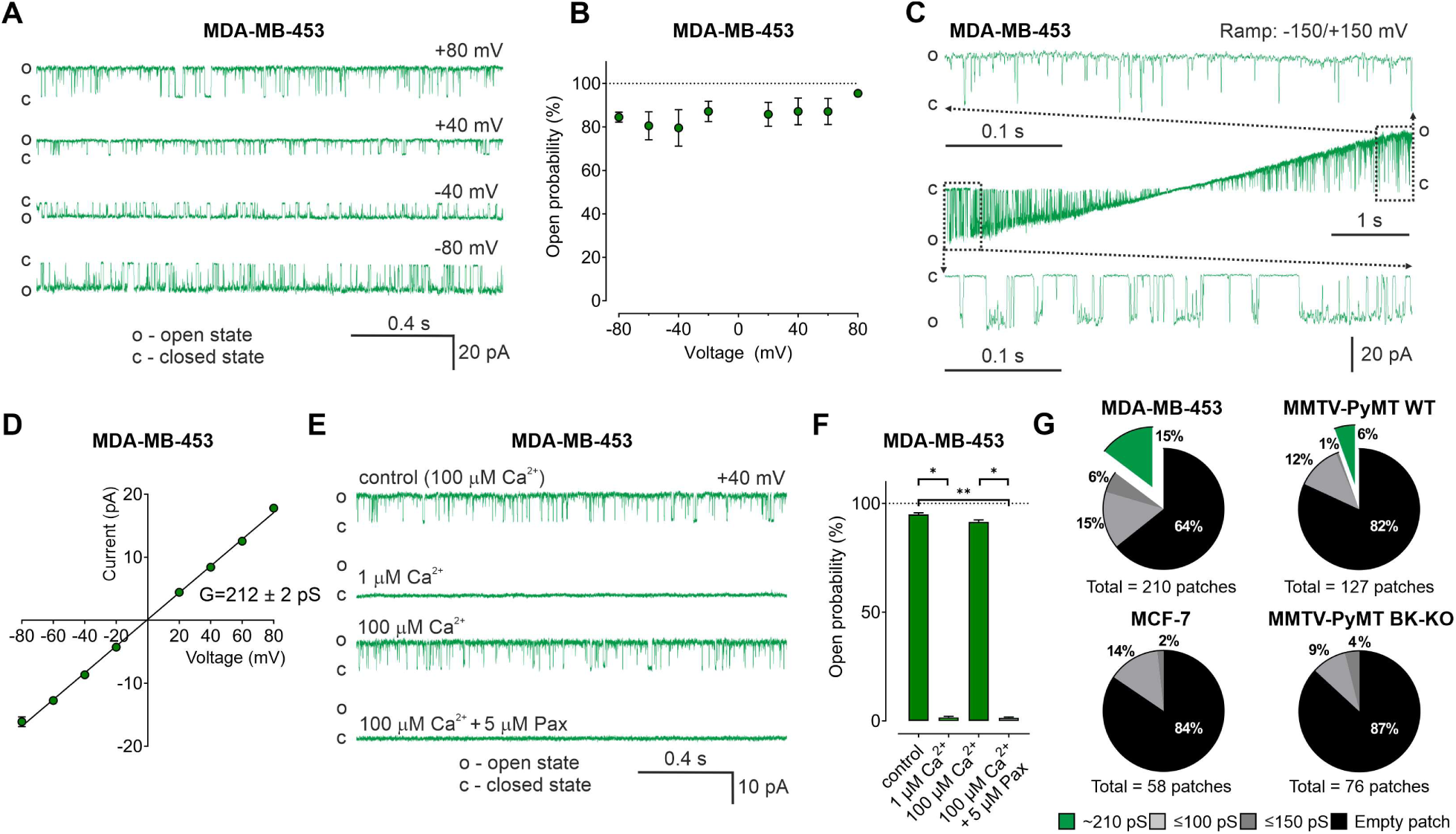
BKCa activity is present in the inner mitochondrial membrane (IMM) of BCCs. (**A**) Representative BKCa single-channel recordings of the IMM of mitoplasts isolated from MDA-MB-453 cells using a symmetric 150/150 mM isotonic KCl solution containing 100 µM Ca^2+^ at voltages ranging from -80 to +80 mV as indicated in the panel. (**B**) Open probability analysis of mitoBKCa at different voltages received from experiments as performed in (**A**). N = 8. (**C**) Single-channel currents of the IMM of mitoplasts isolated from MDA-MB-453 cells recorded using a voltage ramp protocol ranging from −150 to +150 mV. Above and below the ramp are enlarged excerpts of the records shown in rectangles. (**D**) Current-voltage (I-V) plot based on single-channel recordings of MDA-MB-453 cells as performed in **a**, using a symmetric 150/150 mM KCl isotonic solution containing 100 µM Ca^2+^. N = 11. (**E, F**) Representative single-channel recordings of the IMM of mitoplasts isolated from MDA-MB-453 cells (**E**) and corresponding open probabilities at +40 mV in a symmetric 150/150 mM KCl isotonic solution under control conditions (100 μM Ca^2+^), after reducing Ca^2+^ to 1 μM, re-addition of 100 μM Ca^2+^ and finally after application of 5 μM paxilline in the presence of 100 μM Ca^2^. Data in (**F**) show average ± SEM. *p≤0.05, **p≤0.01 using Friedmann test followed by Dunn’s multiple comparison test, n =7. (**G**) Pie chart displaying the percentage of mitoBKCa channel currents (green) possessing a conductance of ∼210 pS, *versus* the total number of patch-clamp experiments performed using mitoplasts isolated from MDA-MB-453 cells (upper left), MMTV-PyMT WT cells (upper right), MCF-7 cells (lower left) and MMTV-PyMT BK-KO cells (lower right). Black segments represent empty patches, bright- and dark grey fraction demonstrate percentage of channels possessing smaller conductances of ≤100 pS and ≤150 pS, respectively. All recordings were low-pass filtered at 1 kHz. “c“ and “o” indicate the closed- and open state of the channel, respectively.

These experiments were further corroborated by immunoblotting experiments using whole-cell lysates and sub-cellular homogenates of different purity. We detected BK_Ca_, the Na^+^/K^+^ ATPase as a marker of the PM, cytochrome c oxidase subunit IV (COXIV) as a marker of mitochondria and TMX1, which localizes to ER membranes in these samples (**Fig. S5D**). A protein band corresponding to BK_Ca_ was identified not only in whole-cell lysates, but also in isolated mitochondria. Importantly, the Na^+^/K^+^ ATPase was absent in the latter protein fraction, confirming the purity of the mitochondrial preparation (**Fig. S5D**). These data confirm the presence of mitoBK_Ca_, potentially BK_Ca_-DEC, in the utilized BK_Ca_ proficient BCCs.

### mitoBK_Ca_ promotes the Warburg effect, triggers cellular O_2_ independency and stimulates BCC proliferation

Based on the observed influence of mitoBK_Ca_ on glycolysis and mitochondrial metabolism, we addressed, whether the channel contributes to the Warburg effect, commonly observed in cancer cells (Bischof et al., 2021; Warburg, 1924). Therefore, we assessed the Warburg index (WI) by investigating the cytosolic lactate concentration ([LAC]_cyto_) over-time using Laconic, a FRET-based lactate sensor (San Martín et al., 2013). [LAC]_cyto_ was followed in response to either inhibiting mitochondrial metabolism by administration of NaN_3_, a complex IV inhibitor, to stop pyruvate consumption (Leary et al., 1998), or upon subsequent inhibition of lactate secretion towards the ECM via monocarboxylate transporter 1 (MCT-1) using BAY-8002 (Quanz et al., 2018) (**Figs. 6A** and **6B**). Indeed, MMTV-PyMT WT cells exhibited an increased WI compared to BK-KO cells under control conditions (**Fig. 6C**), indicating that the presence of BK_Ca_ favors lactate secretion rather than TCA-dependent utilization of pyruvate. Paxilline, but not iberiotoxin treatment reduced the WI in WT cells to the BK-KO level (**Fig. 6C**). The WI profiles of MDA-MB-453 (**Fig. 6D**) and MCF-7 cells (**Fig. 6E**) showed the same sensitivity towards paxilline, while in the latter the expression of BK_Ca_-DEC^RFP^, but not BK_Ca_^RFP^, stimulated the WI. To validate the contribution of the different BK_Ca_ isoforms, endogenous BK_Ca_ transcripts in MMTV-PyMT and MDA-MB-453 cells were targeted using specific siRNAs targeting either the major BK_Ca_ isoforms or specifically the DEC exon of BK_Ca_ (**Fig. S6A**). Cell treatment with the respective siRNAs reduced the expression of BK_Ca_ or BK_Ca_-DEC (**Fig. S6B** and **S6C**). Interestingly, the knockdown of BK_Ca_ interfered with the WI in these cells (**Figs. 6F** and **6G**). This effect was reproduced by specific silencing of the BK_Ca_-DEC isoform (**Figs. 6F** and **6G**), indicating that BK_Ca_-DEC-derived mitoBK_Ca_ channels stimulate the Warburg effect in BCCs.

**Figure 6:**
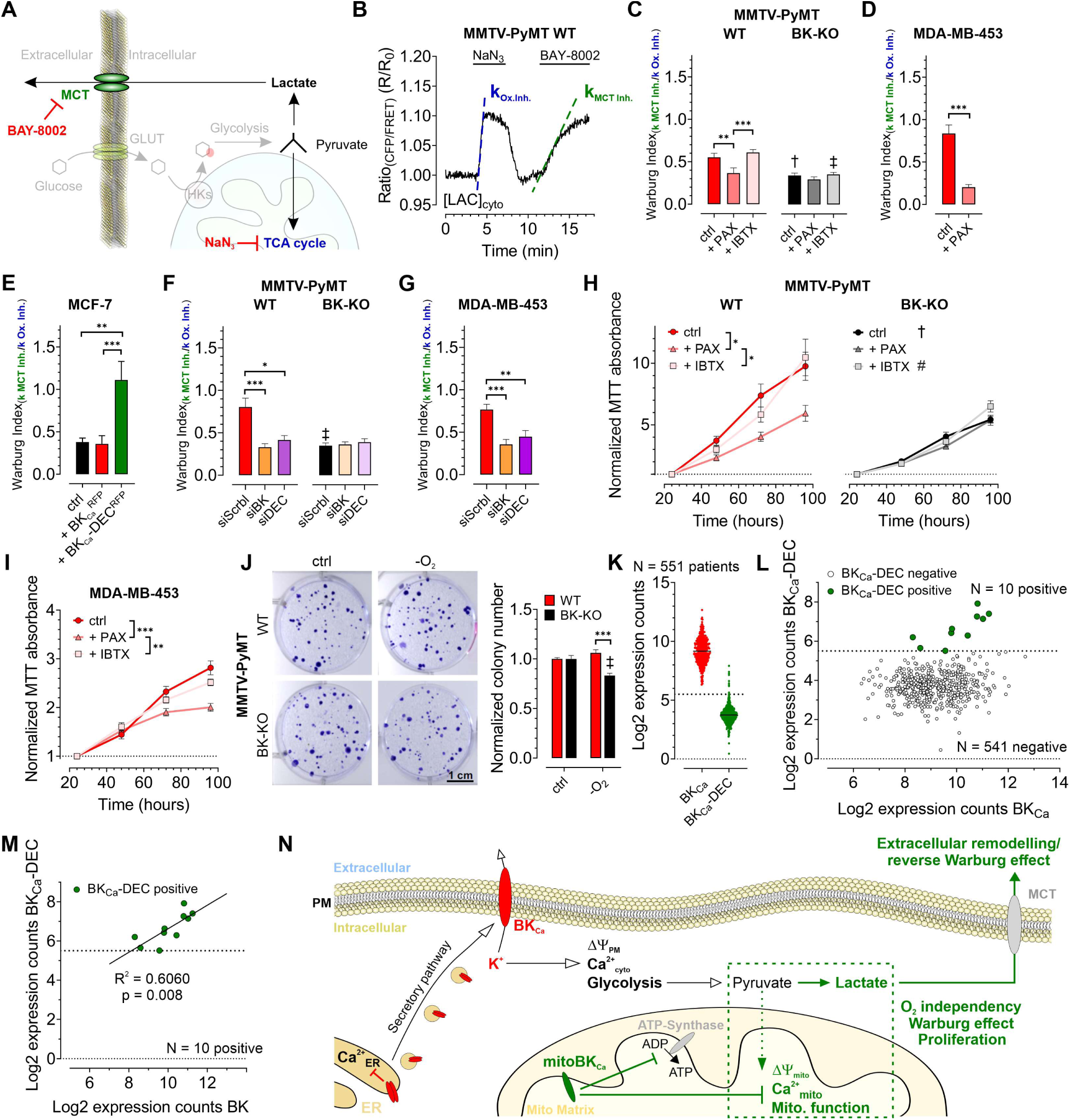
BKCa-DEC expression contributes to the metabolic remodeling and growth of murine and human BCCs and is present in primary tumor samples. (**A**) Schematic representation of the fate of glucose in glycolysis. The tricarboxylic acid (TCA) cycle or lactate secretion via monocarboxylate transporters (MCT) can be inhibited, either using NaN3 or BAY-8002. GLUT: Glucose transporter, HKs: Hexokinases. (**B**) Representative cytosolic lactate concentration ([LAC]cyto) of a MMTV-PyMT WT cell over-time in response to administration or removal of NaN3 and BAY-8002 at time points indicated. Dashed lines indicate slopes taken for assessment of the “Warburg index. (**C – G**) Average Warburg indices ± SEM of MMTV-PyMT WT (**C, F,** left), MMTV-PyMT BK-KO (**C, F**, right), MDA-MB-453 cells (**D, G**) and MCF-7 cells (**E**) calculated from the experiments as shown in (**B**), either under control conditions, in the presence of paxilline or iberiotoxin (**C, D**), upon expression of BKCa^RFP^ or BKCa-DEC^RFP^ (**E**), or upon cell treatment with a scrambled siRNA (siScrbl), or siRNA against a common BKCa sequence targeting all known splice variants (siBK), or a siRNA specifically designed to knockdown BKCa-DEC (siDEC) (**F, G**). (**H, I**) Normalized MTT absorbance over-time of MMTV-PyMT WT (**H**, left) and BK-KO cells (**H**, right), and MDA-MB-453 cells (**I**), either under control conditions, or in the presence of paxilline or iberiotoxin. **J**, Representative images and corresponding statistics of colony formation assays using MMTV-PyMT WT or BK-KO cells in the presence or absence of O2. (**K – N**) mRNA expression of BKCa and BKCa-DEC as performed by Nanostring analysis of 551 BC patient samples. (**K**) Log2 expression counts of BKCa and BKCa-DEC. The threshold for positive expression level was set to log2 = 5.5 (dashed line). (**L**) Log2 expression counts of BKCa-DEC blotted over the log2 expression counts of BKCa. 10 of the 551 patient samples showed expression of BKCa-DEC above the threshold of log2 = 5.5 (dashed line), whereas 541 patient samples were BKCa-DEC negative. (**M**) Correlation of the log2 expression counts of BKCa-DEC positive samples with the log2 expression counts of BKCa in the primary human BC material. (**N**) Summarizing scheme of BKCa in cancer cell homeostasis. N (independent experiments)/ n (cells analyzed) = **C**: 4/26 WT ctrl, 6/28 WT + PAX, 4/39 WT + IBTX, 4/17 BK-KO ctrl, 5/18 BK-KO + PAX, 4/27 BK-KO + IBTX, **D:** 7/29 ctrl, 5/13 + PAX, **E:** 5/27 ctrl, 5/20 + BKCaRFP, 7/26 BKCa-DEC^RFP^, **F:** 5/22 WT siScrbl, 5/28 WT siBK, 4/26 WT siDEC, 5/24 BK-KO siScrbl, 5/24 BK-KO siBK, 5/29 BK-KO siDEC, **G:** 5/21 siScrbl, 5/22 siBK, 5/19 siDEC, **H – J**: 4 for all. *p≤0.05, **p≤0.01, ***p≤0.001, Kruskal-Wallis test followed by Dunn’s MC test (**C, E, F, G, I**), One-Way ANOVA test followed by Tukey’s MC test (**H**) or Mann-Whitney test (**D**). †p≤0.01, ‡p≤0.001 compared to respective WT condition, Unpaired t-test (ctrl, **C, J**) or Mann-Whitney test (+ IBTX, **C, F**).

As glycolytic metabolites and ATP fuel cell proliferation, we investigated the proliferation rates of MMTV-PyMT WT, MMTV-PyMT BK-KO, and MDA-MB-453 cells over-time in the absence or presence of paxilline or iberiotoxin. Using MTT-assay, corroborating previous findings (Mohr et al., 2022), these analyses revealed faster proliferation of WT compared to BK-KO cells under control conditions (**Fig. 6H**). Interestingly, paxilline, but not iberiotoxin treatment, reduced the proliferation of MMTV-PyMT WT to the level of BK_Ca_-deficient cells (**Fig. 6H**). These effects were also observed in MDA-MB-453 cells (**Fig. 6I**). Since the MTT assay is also a redout for mitochondrial function, we additionally performed immunofluorescence-based detection of Ki-67, a frequently used marker of cell proliferation (Dowsett et al., 2011). This approach confirmed the MTT data (**Figs. S6D-F**), establishing intracellular BK_Ca_, possibly mitoBK_Ca_, as key-player in mediating the metabolic rewiring. Next, we studied whether the increased WI affects the proliferation rate of these BCCs at low O_2_ tension (Nazemi and Rainero, 2020). Colony formation assays (CFAs) revealed an increased hypoxic resistance of MMTV-PyMT WT compared to BK-KO cells, as the number and size of colonies lacking BK_Ca_ was reduced upon O_2_-deprivation (**Fig. 6J** and **Fig. S6G**). In sum, these data demonstrate a malignancy-promoting effect of mitoBK_Ca_ in BCC.

### mitoBK_Ca_ is of potential clinical relevance

To determine the clinical relevance of our findings, we investigated whether BK_Ca_-DEC transcripts are present in primary BC tissue. Therefore, the mRNA expression of BK_Ca_ and BK_Ca_-DEC was analyzed by nanostring analysis in bulk tumor biopsies isolated from 551 BC patients. Remarkably, all 551 samples tested positive for BK_Ca_ mRNA expression, with 10 of the samples showing significant expression of BK_Ca_-DEC above the log2 expression threshold of 5.5 (**Figs. 6K** and **6L**). Importantly, if BK_Ca_-DEC was expressed, the expression of BK_Ca_ well correlated with the expression of BK_Ca_-DEC (R^2^ = 0.6060, p = 0.008) (**Fig. 6M**).

In sum, our experiments emphasize the presence of BK_Ca_ in different intracellular organelles including the ER and vesicles of the secretory pathway, yielding its PM localization. At these sites, BK_Ca_ modulates the Ca^2+^ homeostasis and regulates ΔΨ_PM_. K^+^ efflux across the PM may additionally affect glycolysis. Importantly, functionally relevant BK_Ca_ also locates in the IMM of BCCs, promoting, presumably by the K^+^ accumulation in the matrix following channel activation, ΔΨ_mito_ depolarization and consequently ATP-synthase activity in reverse mode as well as a depletion of [Ca^2+^]_mito_. These profound ionic and bioenergetic changes ultimately trigger the proliferation of BCCs in a low oxygen environment, as found in solid tumors (**Fig. 6N**). Taken together, functional expression of mitoBK_Ca_ could possibly denote a prognostic or therapeutic marker for BC patients, and its pharmacologic modulation could represent a novel anti-cancer treatment strategy.

## Discussion

Here, we demonstrate for the first time that BK_Ca_-DEC (mitoBK_Ca_) is functionally expressed in BCCs. BK_Ca_-DEC modulates BCC metabolism, stimulates the Warburg effect, and accelerates cell proliferation rates in the presence and absence of O_2_. These tumor- and malignancy-promoting effects were sensitive to BK_Ca_ inhibition using the cell-permeable BK_Ca_ inhibitor paxilline (Zhou and Lingle, 2014), but not the cell-impermeable blocker iberiotoxin (Candia et al., 1992), indicating that intracellular BK_Ca_, presumably mitoBK_Ca_, mediates malignant BCC behavior and tumor development.

In line with recent single-cell RNA sequencing data of 26 primary breast tumors (S. Z. Wu et al., 2021), we found high transcript levels for BK_Ca_ throughout the analyzed BC samples, in addition to BK_Ca_-DEC, albeit in a much smaller subset of BC. The analyzed BC samples were, however, all positive for hormone receptors. Whether BK_Ca_ expression is different in hormone receptor negative specimens hence needs to be further investigated. Further, we confirmed functional BK_Ca_ expression in the PM of MMTV-PyMT WT and MDA-MB-453 cells, while MMTV-PyMT BK-KO and MCF-7 cells showed no or very low PM BK_Ca_ currents. Therefore, MCF-7 cells represented a suitable model to investigate the effects of BK_Ca_ over-/expression on the metabolic homeostasis of human BCC. If the low BK_Ca_ expression correlates with hormonal receptor status or alternatively with human epidermal growth factor 2 (HER2) expression levels must be clarified by future studies. Interestingly, however, recent data have shown that BK_Ca_ is overexpressed in triple-negative BC, a fact that led the authors to draw similar conclusions to ours, that BK_Ca_ may represent a novel anti-cancer treatment strategy for selected BC patients (Sizemore et al., 2020). Nevertheless, similarly to cardiac myocytes (CM), expression of BK_Ca_-DEC yielded mitochondrial localization of BK_Ca_-DEC, although its abundance in the IMM appeared less pronounced compared to CM (Singh et al., 2013). While a direct comparison is difficult as plasmid transfections may have caused unexpected effects, our finding that BK_Ca_-DEC^RFP^ caused significantly reduced currents across the PM compared to BK_Ca_^RFP^ putatively confirm its increased intracellular abundance.

Based on the potential impact of BK_Ca_ on the cellular ΔΨ_PM_ and ion balance (Burgstaller et al., 2022a), we conducted an in-depth investigation of the (sub)cellular Ca^2+^ homeostasis. Across the BCCs examined, we observed that functional BK_Ca_ expression modulated [Ca^2+^]_cyto_ dynamics. These alterations showed, however, differential sensitivities to BK_Ca_ inhibitors in MMTV-PyMT WT and MDA-MB-453 cells. While basal [Ca^2+^]_cyto_ in MDA-MB-453 cells was reduced by both, paxilline and iberiotoxin treatment, MMTV-PyMT WT cells were only sensitive to paxilline. Examination of [Ca^2+^]_ER_ confirmed the results from [Ca^2+^]_cyto_, as [Ca^2+^]_ER_ levels increased with paxilline and iberiotoxin in MDA-MB-453, but only upon paxilline exposure in MMTV-PyMT WT cells, suggesting differential effects of intracellular and PM-localized BK_Ca_ on Ca^2+^ handling in these cells. Interestingly, in MCF-7 cells, both BK_Ca_ splice variants mediated the opposing effects on the basal [Ca^2+^]_cyto_ and [Ca^2+^]_ER_ levels, which either in- or decreased, respectively, upon transient BK_Ca_^RFP^ or BK_Ca_-DEC^RFP^ expression. This is expected as the transitory hyperpolarization and the efflux of K^+^ due to the opening of PM BK_Ca_ provides the driving force for Ca^2+^ entry into cytoplasm, while a BK_Ca_-mediated K^+^ increase within the ER, presumably through PM-directed channels crossing the ER membrane in the secretory pathway, would oppose the Ca^2+^ refiling (Burgstaller et al., 2022a). Moreover, these results are in line with the higher proliferative capability of BK_Ca_ proficient BCCs due to the manifold roles of Ca^2+^ as second messenger (Burgstaller et al., 2022a). Importantly, however, [Ca^2+^]_mito_ of BK_Ca_ proficient MMTV-PyMT WT and MDA-MB-453 cells was exclusively sensitive for paxilline, and it was specifically affected by the expression of BK_Ca_-DEC^RFP^, but not by BK_Ca_^RFP^, in MCF-7 cells, suggesting that this Ca^2+^ pool is exclusively controlled by intracellular BK_Ca_.

Subcellular Ca^2+^ alterations could reportedly alter cellular bioenergetics, as Ca^2+^ directly regulates metabolic enzymes and activities (Rossi et al., 2019). Indeed, extracellular flux analysis, LC-MS-based metabolomics and fluorescence-based live-cell imaging confirmed, that the observed alteration in sub-cellular Ca^2+^ homeostasis caused by BK_Ca_, especially mitoBK_Ca_, has severe effects on cell metabolism. Our data further emphasize, that the presence of mitoBK_Ca_, as confirmed by mitoplast patch-clamp and Western blot analysis of isolated mitochondria, depolarizes BCC mitochondria, which is in line with a previous study showing an impact of mitoBK_Ca_ activation on ΔΨ_mito_ (Kicinska et al., 2016). mitoBKCa-dependent depolarization of ΔΨ_mito_ in turn triggers cellular glucose dependency and reverses the activity of the mitochondrial ATP-synthase to consume ATP for restoring ΔΨ_mito_. Finally, BK_Ca_-DEC-derived mitoBK_Ca_ channels promote the Warburg effect and ultimately stimulated proliferation rates of BCCs. Our data fit earlier findings from glioma cells, where an O_2_-sensitivity of mitoBK_Ca_ was observed, which probably increased the hypoxic resistance of these cancer cells (Gu et al., 2014). It may seem contradictory that mitoBK_Ca_ is highly expressed in CM, which are among one of the most oxidative cell types known. It is, however, important to mention, that CM, under physiologic conditions, do not show a metabolic Warburg setting, which is a common phenomenon of cancer cells propelling their O_2_ independency, due to the hypoxic microenvironment. Moreover, it was demonstrated recently that the absence of (mito)BK_Ca_ does not alter physiologic cardiac function. Only upon induction of ischemia and reperfusion injury, a lack of (mito)BK_Ca_ promoted the susceptibility of the heart to cell death, resulting in increased infarction size (Frankenreiter et al., 2017). Hence, it can be concluded that (mito)BK_Ca_ only played a role under conditions where mitochondria are, due to the absence of O_2_, not properly functioning. Taking this view into account, the results derived from CM are consistent with our findings in BCCs, as (mito)BK_Ca_ mediates the resistance to hypoxic stress to BCC.

Excessive production and release of lactate, a hallmark of the Warburg effect, leads to extracellular acidification, subsequently creating a microenvironment that promotes tumorigenesis and metastasis as well as the resistance to anti-tumor immune responses and therapy (de la Cruz-López et al., 2019; Nazemi and Rainero, 2020; P. Wu et al., 2021). Extracellular K^+^ [K^+^]_ex_, in turn, accumulating within the necrotic core of solid tumors, was shown to interfere with effector T-cell function triggering immune escape of cancer cells (Eil et al., 2016). To elucidate whether mitoBK_Ca_ directly contributes to lactate-induced tumor aggressiveness or [K^+^]_ex_, live-cell imaging of extracellular metabolites in 3D BCC models should be applied in future investigations (Burgstaller et al., 2022b, 2021b).

Finally, our results demonstrate for the first time that BK_Ca_-DEC transcripts are present in human BC biopsies. Although only a small proportion of patients was positive for BK_Ca_-DEC expression, this finding could be of considerable clinical relevance considering the link between mitoBK_Ca_ function and BCC metabolism. Importantly, the design of our study likely underestimates the incidental number of BK_Ca_-DEC positive BC, as i.) only hormone-receptor positive BC specimens were included, and ii.) bulk-tumor mRNA was analyzed, hampering the detection of low-abundant or tightly regulated transcripts against a strong background of non-cancer cells present in bulk tumor tissues. Finally, due to the small number of positive hits (N=10 positive *versus* N=541 BK_Ca_-DEC negative specimens) and the lack of (sufficient) follow-up information in some of these cases, we are currently unable to correlate BK_Ca_-DEC expression with, for example, treatment response or survival. Thus, future studies are warranted to show how the tumor’s BK_Ca_-DEC status can help to predict or therapeutically improve standard chemo-endocrine treatments. The rather low abundance of BK_Ca_-DEC in the clinical samples, however, is in agreement with our mitoplast patch clamp experiments with mitoBK_Ca_-mediated K^+^ currents being detected at frequencies between 6 to 15%, suggesting that either a small proportion of mitochondria express functional mitoBK_Ca_, or that the abundance of the channel per mitochondrion is low, requiring sensitive mechanistic approaches for its detection.

In summary, our data highlight a potentially druggable mitoBK_Ca_ isoform in BCCs, whose molecular entity is mainly formed by the *Kcnma1* encoded BK_Ca_-DEC splice variant. This channel promotes metabolic alterations in cancer cells, even under low-oxygen conditions, which may ultimately be of clinical interest for new anti-cancer therapies.

## Material and Methods

### Buffers and solutions

If not otherwise stated, all chemicals were purchased from Carl Roth GmbH, Karlsruhe, Germany.

Buffers used in this study comprised the following:

Physiologic buffer for single cell live recordings contained (in mM): 138 NaCl, 5 KCl, 2 CaCl_2_, 1 MgCl_2_, 2 glucose, 10 HEPES, pH set to 7.4 with NaOH. No glucose was added during glucose removal experiments, while glucose was increased to 25 mM to investigate glucose dependency of the mitochondrial membrane potential. 10 mM CaCl_2_ instead of 2 mM CaCl_2_ were added to obtain 10Ca buffer. 0.1 mM Ethylene glycol bis(2-aminoethylether)-N, N, N’, N’-tetra acetic acid (EGTA) (Sigma Aldrich Chemie GmbH, Taufkirchen, Germany) instead of 2.0 mM CaCl_2_ was added to obtain Ca^2+^ free buffer. For 0 mM K^+^ buffer, 5 mM KCl was replaced by 5 mM NaCl. For 300 mM K^+^ buffer, 300 mM KCl was added instead of 5 mM K^+^ and addition of NaCl was omitted. The following compounds were added to yield the following final concentration: 3 µM Oligomycin-A, 200 nM Carbonyl cyanide-p-trifluoromethoxyphenylhydrazone (FCCP), 5 µM paxilline (all from (Santa Cruz Biotechnology, Dallas, USA), 30 nM iberiotoxin (Selleckchem, Planegg, Germany), 5 mM NaN_3_, 3 µM BAY-8002, 15 µM 2,5-Di-(t-butyl)-1,4-hydroquinone (BHQ) (all from Sigma Aldrich Chemie GmbH), 100 µM adenosine-5’-triphosphate (ATP), 5 µM ionomycin (Alomone Labs, Jerusalem, Israel), 15 µM gramicidin (Sigma Aldrich Chemie GmbH). For H_2_O insoluble compounds, final DMSO concentration in the buffer did not exceed 0.1%.

Intracellular buffer used for whole-cell patch-clamp experiments contained (in mM): 130 K-Gluconate, 5 KCl, 2 Mg-ATP, 0.1 CaCl_2_, 0.2 Na_2_-GTP, 0.6 EGTA, 5 HEPES, pH = 7.2 with KOH.

Cell equilibration buffer fluorescence microscopy-based contained (in mM): 135 NaCl, 5 KCl, 2 CaCl_2_, 1 MgCl_2_, 2.6 NaHCO_3_, 0.44 KH_2_PO_4_, 0.34 Na_2_HPO_4_, 10 glucose, 10 HEPES, 2 GlutaMAX, 1 sodium pyruvate, with 1x MEM amino acids and 1x MEM vitamins added (both Thermo Fisher Scientific). pH was adjusted to 7.4 using NaOH.

Buffers used for mitochondrial isolation, mitoplast preparation, and single channel patch-clamp comprised the following: The preparation solution contained (in mM): 250 sucrose, 5 HEPES, pH = 7.2. The mitochondrial storage buffer contained (in mM): 150 KCl, 0.1 CaCl_2_, 20 HEPES, pH = 7.2. The hypotonic buffer contained (in mM) 0.1 CaCl_2_, 5 HEPES, pH = 7.2. The hypertonic buffer contained (in mM): 750 KCl, 0.1 CaCl_2_, 30 HEPES, pH 7.2. Low-Ca^2+^ solution (1 μM Ca^2+^) contained (in mM): 150 KCl, 1 EGTA, 0.752 CaCl2, 10 HEPES, pH = 7.2.

### Cell culture and transfection

Mouse mammary tumor virus polyoma middle T antigen (MMTV-PyMT) cells were isolated from tumors of MMTV-PyMT transgenic FVB/N WT or BK-KO mice. Tumor growth *in vivo* and biopsies were authorized by the local ethics *Committee for Animal Research* (Regierungspräsidium Tuebingen, Germany), and were performed in accordance with the *German Animal Welfare Act*. Animals were kept on a 12-hour light/ dark cycle under temperature- and humidity-controlled conditions with unlimited access to food (Altromin, Lage, Germany) and water. MMTV-PyMT cells used in this study were isolated from 3 – 7 different female breast-cancer bearing WT and 3 – 4 different female breast-cancer bearing BK-KO animals at an age of ∼ 12 – 14 weeks. Upon dissection, tumors were carefully minced into pieces using atraumatic forceps, lysed by 1 mg mL^-1^ Collagenase-D (Roche, Basel, Switzerland) for 10 min, and cultured as follows: Cells were grown in modified improved minimal essential medium (IMEM) supplemented with 5% fetal bovine serum (FBS), 1 mM sodium pyruvate and 100 U mL^−1^ penicillin and 100 µg mL^−1^ streptomycin (all purchased from Thermo Fisher Scientific) at 37°C and 5% CO_2_ in a humidified incubator. Fibroblasts were removed by exposure of the cultures to 0.25% trypsin-EDTA in PBS (Thermo Fisher Scientific) and short incubation at 37°C (∼ 1 minute). After gently tapping the plate, trypsin-EDTA with detached fibroblasts was removed and cells were further cultured in supplemented modified IMEM at 37°C and 5% CO_2_ until subculturing.

MCF-7 and MDA-MB-453 cells were purchased from the Global Bioresource Center (ATCC). Cells were cultivated in Dulbecco’s modified eagle’s medium (DMEM) supplemented with 10% FBS, 1 mM sodium pyruvate and 100 U mL^−1^ penicillin and 100 µg mL^−1^ streptomycin (Thermo Fisher Scientific) at 37°C and 5% CO_2_ in a humidified incubator.

Subculturing of cells was performed when cells reached a confluency of 80 – 90%. Therefore, cell culture medium was removed, cells were washed 1x with PBS, and trypsin-EDTA at a final concentration of 0.25% trypsin-EDTA in PBS was added. Subsequently, cells were incubated at 37°C and 5% CO_2_ in a humidified incubator until cell detachment occurred (∼ 2-5 minutes). Trypsinization was stopped by adding supplemented cell culture media and cells were pelleted at 300xg for 5 minutes. The supernatant was removed, and cells were seeded to new cell culture dishes as required. For fluorescence microscopic live-cell imaging experiments, cells were either seeded in 6-well plates containing 1.5 H 30 mm circular glass coverslips (Paul Marienfeld GmbH, Lauda-Königshofen, Germany). All other vessels and serological pipettes used for cell culture were ordered from Corning (Kaiserslautern, Germany).

Transfection of cells was performed according to manufacturer’s instructions when cells showed a confluency of ∼70%, either using PolyJET DNA transfection reagent (SignaGen Laboratories, Maryland, USA) for plasmid DNA transfection or Lipofectamine 2000 (Thermo Fisher Scientific) for siRNA transfection or co-transfection of siRNA with plasmid DNA. Plasmid DNA amount was reduced to 1/3 for transfection of mitochondrial-targeted probes to ensure proper mitochondrial localization of the probes. Plasmid transfections were performed 16 hours, siRNA transfections were performed 48 hours before the experiments. Paxilline or iberiotoxin were added 12 hours prior to the experiments to the respective cell culture medium. DMSO served as a control.

### Whole-cell patch-clamp

For whole-cell patch-clamp experiments, 30,000 cells were seeded on the day before the experiment in 35 mm glass bottom µDishes (ibidi GmbH, Graefelfing, Germany) and cultivated in the respective supplemented cell culture medium over-night at 37°C and 5% CO_2_ in a humidified incubator. The next day, cell culture medium was removed, and cells were washed 2x and maintained in prewarmed physiologic buffer. Subsequently, recordings were performed using borosilicate glass capillaries (0.86x1.5x100mm) (Science Products GmbH, Hofheim am Taunus, Germany), with a resistance of 4-6 MW, which were pulled using a model P-1000 flaming/ brown micropipette puller (Sutter Instruments, California, USA) and filled with intracellular buffer. A MP-225 micromanipulator served for pipette control (Sutter Instruments). Recordings were performed in whole-cell mode. Currents were evoked by 15 voltage square pulses (300 ms each) from the holding potential of -70 mV to voltages between -100 mV and +180 mV delivered in 20 mV increments. For amplifier control (EPC 10) and data acquisition, Patchmaster software (HEKA Elektronik GmbH, Lambrecht, Germany) was used. Voltages were corrected offline for the capacity. Data analysis was performed using Fitmaster software (HEKA Elektronik GmbH), Nest-o-Patch software (http://sourceforge.net/projects/nestopatch, written by Dr. V Nesterov), and Microsoft Excel (Microsoft, Washington, USA).

### Cloning and plasmid preparation

Cloning was performed using conventional PCR-, restriction- and ligation-based procedures. BK_Ca_-DEC was a gift from Michael J. Shipston and was N-terminally attached to an RFP (BK_Ca_-DEC^RFP^) using KpnI and BamHI restriction sites after PCR amplification (NEB Q5 High-Fidelity DNA-Polymerase, New England Biolabs (NEB), Ipswich, USA). For generation of BK_Ca_^RFP^, a PCR amplification of BK_Ca_-DEC was performed, where the reverse primer omitted the last amino acids including the 50 amino acids encoding the DEC exon. Mitochondrial targeted TagRFP (mtRFP) was generated by fusing a double repeat of COX8 pre-sequence N-terminally to TagRFP. RFP-GPI was generated by fusing the membrane leading sequence (MLS) and the GPI-anchor signal of cadherin 13 N- and C-terminally to TagRFP, respectively. After PCR reactions, the DNA fragments were purified from gel electrophoresis using the Monarch DNA gel extraction kit (NEB), fragments and destination plasmid were digested using the respective restriction enzymes (NEB) and ligation (T4 DNA Ligase, NEB) and transformation (chemically competent NEB 5-alpha *E. coli*) were performed according to manufacturer’s instructions. Plasmids were verified by sequencing (Microsynth AG, Balgach, Switzerland). DNA maxipreps were performed using the Nucleobond Xtra Maxi kit (Macherey Nagel GmbH & Co. KG, Düren, Germany). Purified DNA was stored at 4°C.

### Confocal imaging

For confocal imaging of mtRFP, RFP-GPI, BK_Ca_^RFP^ or BK_Ca_-DEC^RFP^ colocalization with mitochondria, MCF-7 cells were seeded on circular 30 mm glass coverslips (Marienfeld GmbH) in 6-well plates (Corning). Cells were transfected using PolyJet transfection reagent according to the manufacturer’s instructions. 16 hours after transfection medium was exchanged for fresh cell culture medium and cells were further cultivated for 24 hours. Subsequently, the medium was exchanged for cell equilibration buffer containing MitoGREEN (PromoCELL GmbH, Heidelberg, Germany) at a final concentration of 3 µM and cells were incubated at room temperature for 30 minutes. Subsequently, cells were washed 2x with physiologic buffer, and cells were analysed using confocal fluorescence microscopy Imaging was performed using a Zeiss LSM 980 equipped with an Airyscan 2 detector. A Zeiss C Plan-Apochromat 63x/1,4 Oil DIC M27 objective was used for all images. The ZEN 3.7 software (blue edition) was used for image acquisition and super resolution images were processed using the ZEN Airyscan module (Carl Zeiss AG). Tag-RFPs were excited at 561 nm and detected at 380-735 nm. MitoGREEN was excited at 488 nm and detected at 495-550 nm. Image analysis was performed using the colocalization test in ImageJ with Fay randomization after cell selection by ROIs.

### Fluorescence live-cell imaging

Cells were either analyzed using a Zeiss AXIO Observer Z1 or a Zeiss Axiovert 200m microscope (Carl Zeiss AG, Oberkochen, Germany). The Zeiss AXIO Observer Z1 was connected to a LEDHub high-power LED light engine (OMICRON Laserage, Rodgau-Dudenhofen, Germany) and equipped with a EC Plan-Neofluar 40x/1.3 Oil DIC M27 objective (Carl Zeiss AG), an Optosplit II emission image splitter (Cairn Research Ltd, Faversham, UK), and a pco.panda 4.2 bi sCMOS camera (Excelitas PCO GmbH, Kelheim, Germany). The microscope possessed a BioPrecision2 automatic XY-Table (Ludl Electronic Products, Ltd., New York, USA). Optical filters included a 459/526/596 dichroic mirror and a 475/543/702 emission filter for FRET- and TMRM-based measurements, and a 409/493/573/652 dichroic mirror combined with a 514/605/730 emission filter for Dibac4(3), Fura-2 and 2-NBDG based measurements (all purchased from AHF Analysentechnik, Tuebingen, Germany). The Optosplit II emission image splitter was equipped with a T505lpxr long-pass filter (AHF Analysentechnik). The LEDHub high-power LED light engine was equipped with a 340 nm, 385 nm, 455 nm, 470 nm and 505-600 nm LED, followed by the following emission filters, respectively: 340x, 380x, 427/10, 473/10 and 510/10 or 575/15 (AHF Analysentechnik). The Zeiss Axiovert 200m microscope was connected to a pe340^fura^ light source (CoolLED, Andover, UK), an Optosplit II emission image splitter (Cairn Research Ltd.) and a pco.panda 4.2 sCMOS camera (Excelitas PCO GmbH) and equipped with 340/26, 380/14 and switchable 427/10, 485/20 or 575/15 excitation filters (AHF Analysentechnik) in the light source, respectively, a 40x Fluar 1.30 oil immersion objective (Carl Zeiss AG), a 459/526/596 or 515LP dichroic mirror and a 475/543/702 or 525/15 emission filter (AHF Analysentechnik) in the microscope, and a T505lpxr (AHF Analysentechnik) in the Optosplit II. Image acquisition and control of both microscopes was performed using VisiView software (Visitron Systems GmbH, Puchheim, Germany). Perfusion of cells was performed using a PC30 perfusion chamber connected to a gravity-based perfusion system (NGFI GmbH, Graz, Austria) and a vacuum pump.

### Fura-2 based Ca^2+^ measurements

For fura-2 based Ca^2+^ measurements, cells were taken from the humidified incubator at 37°C and 5% CO_2_, washed 1x with cell equilibration buffer and loaded with fura-2 AM (Biomol GmbH, Hamburg, Germany) at a final concentration of 3.3 µM in cell equilibration buffer for 45 minutes at room temperature. Subsequently, cells were washed 2x with cell equilibration buffer and stored in equilibration buffer for additional 30 minutes prior to the measurements. Paxilline or iberiotoxin treatment of the cells was performed 12 hours prior to the measurements and both inhibitors remained present during the fura-2 loading procedure and the measurement at concentrations of 5 µM and 30 nM, respectively. Imaging experiments were either performed on the Zeiss AXIO Observer Z1 or the Zeiss Axiovert 200m microscope (Carl Zeiss AG) in physiologic buffer using alternate excitations at 340 nm and 380 nm. Emissions were captured at roughly 514 nm (Zeiss AXIO Observer Z1) or 525 nm (Zeiss Axiovert 200m). To evoke intracellular Ca^2+^ signals, cells were perfused with physiologic buffer containing ATP (Carl Roth GmbH) at a final concentration of 100 µM.

### Ca^2+^ and K^+^ calibrations

Calibrations of [Ca^2+^]_cyto_ and [K^+^]_cyto_ were performed by initial superfusion of cells in physiologic buffer. Subsequently, buffer was exchanged to Ca^2+^ free buffer containing 5 µM of ionomycin (Alomone Labs), and subsequent switching to 10Ca buffer for Fura-2 saturation, or 0 mM K^+^ buffer containing 15 µM gramicidin, followed by switching to 300 mM K^+^ buffer for saturation of NES lc-LysM GEPII 1.0. For calculation of the [Ca^2+^]_cyto_ and [K^+^]_cyto_ in nM and mM, respectively, the following formula was used:

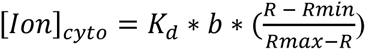

### Genetically encoded sensor-based measurements

Cells were grown on 1.5 H 30 mm circular glass coverslips (Paul Marienfeld GmbH) for perfusion-based experiments. Cells were taken from the incubator, cell culture medium was removed, and cells were equilibrated for at least 30 minutes in cell equilibration buffer in ambient environment. Sensor plasmids used in this study comprised the following: D1ER (Palmer et al., 2004) (Addgene plasmid #36325) for measurement of [Ca^2+^]_ER_ and 4mtD3cpV (Palmer et al., 2006) (Addgene plasmid #36324) for measurement of [Ca^2+^]_mito_ were a gift from Amy Palmer & Roger Tsien, mtAT1.03 (Imamura et al., 2009) for measurement of [ATP]_mito_ was a gift from Hiromi Imamura, Laconic (San Martín et al., 2013) (Addgene plasmid #44238) for measurement of [LAC]_cyto_ and assessment of the Warburg index was a gift from Luis Felipe Barros, mito-Hyper3 (Bilan et al., 2013) was a gift from Markus Waldeck-Weiermair and NES lc-LysM GEPII 1.0 (Bischof et al., 2017) was a gift from Roland Malli.. Experiments were either performed on the AXIO Observer Z1 or the Axiovert 200m microscope (Carl Zeiss AG). For experiments where paxilline (5 µM) or iberiotoxin (30 nM) were used, these compounds were added with the media change prior to the transfection with the PolyJet transfection reagent (SignaGen Laboratories) approximately 12 hours prior to the experiments. The compounds remained present throughout the experiment. FRET-based sensors were excited at 427/10 nm and emissions were collected simultaneously at roughly 475 and 543 nm. mito-Hyper3 was excited at 427/10 and 510/10 nm. Calibration of NES lc-LysM GEPII 1.0 was performed by superfusing cells with 15 µM Gramicidin (Sigma Aldrich Chemie GmbH) either in the absence of K^+^ (0 mM K^+^ buffer) or in the presence of 300 mM K^+^. (300 mM K^+^ buffer).

### Extracellular flux analysis

Assessment of extracellular acidification rate (ECAR) and oxygen consumption rate (OCR) was performed using a Seahorse XFe24 analyzer (Agilent, Santa Clara, USA). The day before the assay, Seahorse XF24 cell culture microplates (Agilent) were coated with 0.5 mg mL^-1^ poly-L-lysine (Sigma Aldrich Chemie GmbH) for 30 minutes at 37°C, washed 2x with PBS followed by cell seeding of 50,000 MMTV-PyMT WT, 50,000 MMTV-PyMT BK-KO or 100,000 MDA-MB-453 cells per well of the 24-well plate in a final volume of 250 µL, either in the presence or absence of 5 µM paxilline, 30 nM iberiotoxin or an equivalent amount of DMSO. 4 wells contained medium without cells and served as blank. Cells were cultivated over-night at 37°C and 5% CO_2_ in a humidified incubator. The Seahorse XFe24 sensor cartridges were equilibrated over-night at 37°C in Seahorse XF Calibrant solution according to manufacturer’s instructions. The next day, cells were washed using Seahorse XF DMEM medium, pH 7.4 additionally supplemented with 5.5 mM glucose, 2 mM GlutaMAX and 1 mM sodium pyruvate (Thermo Fisher Scientific), with or without paxilline, iberiotoxin or DMSO according to manufacturer’s instructions. With the last washing step, a final volume of 500 µL was adjusted per well and cells were incubated at 37°C in the absence of CO_2_ for 30 minutes. Meanwhile the Seahorse XFe24 sensor cartridge was loaded with the following compounds: 55 µL of 20 µM Oligomycin-A, 62 µL of 3 µM FCCP and 69 µL of 25 µM Antimycin-A (Santa Cruz Biotechnology), yielding final concentrations of 2 µM, 300 nM and 2.5 µM upon injection, respectively. For analysis, ECAR and OCR rates were normalized for the protein concentration (µg) per well, which was assessed using the Pierce BCA protein assay kit (Thermo Fisher Scientific) according to manufacturer’s instructions. Absorbance was measured at 540 nm using a TECAN multiplate reader (TECAN Group Ltd., Männedorf, Switzerland) and concentrations were assessed using a calibration curve.

### LC-MS-based metabolomics

Formic acid, acetic acid, acetonitrile and methanol of Ultra LC-MS grade were supplied by Carl Roth (Karlsruhe, Germany). Ammonium hydroxide solution (Suprapur® quality 28.0 - 30.0% NH_3_ basis) was purchased from Sigma-Aldrich (Merck, Taufkirchen, Germany). Deionized water was purified by a Purelab ultrapurification system (ELGA LabWater, Celle, Germany). Uniformly (U-) ^13^C-labeled yeast extract of more than 2 × 109 Pichia pastoris cells (∼15 mg; strain CBS 7435) was obtained from ISOtopic Solutions (Vienna, Austria). All standards used were purchased from Sigma-Aldrich Chemie GmbH. Stock solutions of the individual calibrants were prepared at concentrations of 1 mg mL^-1^ and used for further dilution. The individual stocks were stored at -80°C until use.

Targeted LC-MS analysis was performed using an Agilent 1290 Infinity II series UHPLC system from Agilent Technologies (Waldbronn, Germany) equipped with a binary pump, autosampler, thermostated column compartment and a QTrap 4500 mass spectrometer with a TurboIonSpray Source from SCIEX (Ontario, Canada). The samples were filled into homogenization tubes. Internal standards were added prior to sample preparation. For the extraction of the analytes, 1 mL of the ice-cold extraction solvent (50% methanol and 50% water) and 0.15 g of zirconia/glass beads were added to the cell pellets (which were slowly thawed on ice). The samples were homogenized (6,800 rpm, 1 min at 4 °C, 10 × 10 s, pause 30 s) with Cryolys Evolution using dry ice cooling (Bertin Technologies, France). The samples were then spun down for 5 min (16,100xg at 4 °C). The supernatant was carefully removed, transferred into fresh tubes and evaporated to dryness overnight under nitrogen using a high-performance evaporator (Genevac EZ-2) (Genevac, Ipswich, UK). The dry residue of the extract was reconstituted in 10 µL water and 90 µL acetonitrile, followed by 3 cycles of vortexing and sonication (30 seconds each). The samples were centrifuged at 18928xg for 5 min and the supernatant was used for further analysis.

Chromatographic separation was performed on a Waters (Eschborn, Germany) Premier BEH Amide column (150 x 2.1 mm, 1.7 µm). For metabolite analysis in ESI^+^ mode, mobile phases A and B were adjusted to a pH of 3.5 with formic acid and consisted of 20 mM ammonium formate in water and acetonitrile, respectively. In ESI-, the chromatographic conditions differed. Mobile phase A and B were adjusted to a pH of 7.5 with acetic acid and consisted of 20 mM ammonium acetate in water and acetonitrile, respectively. The gradient elution profile was the same for both positive and negative ionization modes (0.0 min, 100% B; 13 min 70% B; 15 min 70% B; 15.1 min 100% B; 20 min 100% B) and was carried out at a flow rate of 0.25 mL min^-1^ and a constant column temperature of 35 °C. The injection volume was 5 µL. The autosampler was kept at 4°C. Ion source parameters were as follows: nebulizer gas (GS1, zero grade air) 50 psi, heater gas (GS2, zero grade air) 30 psi, curtain gas (CUR, nitrogen) 30 psi, source temperature (TEM) 450 °C, ion source voltage +5,500 V (positive mode) and - 4,500 V (negative mode). Due to the large number of transitions monitored simultaneously, the Scheduled-MRM function was enabled. A window of 30 seconds was set around the designated metabolite-specific retention time and the total cycle time was 1 s. Blank solvents (mobile phase A and B) followed by QCs were injected in the beginning of the chromatographic batch to ensure proper column and system equilibration.

### Plasma- and mitochondrial membrane potential measurements

ΔΨ_PM_ was determined using Bis-(1,3-dibutylbarbituric acid)trimethine oxonol (Dibac4(3)) (Thermo Fisher Scientific). Cells were cultivated on 30 mm glass coverslips. On the day of analysis, cells were equilibrated in cell equilibration buffer containing Dibac4(3) at a concentration of 0.25 µg mL^-1^ for 30 minutes and subsequently analyzed by fluorescence microscopy using 485/20 excitation at the Zeiss Axiovert 200m microscope. Emission was captured at roughly 525 nm.

ΔΨ_mito_ was assessed using Tetramethylrhodamin-Methylester (TMRM) (Thermo Fisher Scientific). After cell cultivation on 30 mm glass coverslips in the presence or absence of paxilline, iberiotoxin or DMSO for 12 hours, the cell culture medium was replaced with cell equilibration buffer containing 200 nM TMRM and cells were incubated in ambient environment for 30 minutes. Subsequently, cells were washed with physiologic buffer containing 2 mM or 25 mM glucose and 200 nM TMRM, and cells were equilibrated for further 30 minutes. The glass coverslips were transferred to the PC30 perfusion chamber and experiments were started using the gravity-based perfusion system (NGFI GmbH). Paxilline, iberiotoxin or DMSO (control) and 200 nM TMRM remained present throughout the experiment. Mitochondrial depolarization was induced by the perfusion of a buffer containing 200 nM FCCP. For analysis, the fluorescence emission at ≥600 nm upon excitation at 575/15 nm of a region of interest (ROI) above mitochondria was divided by a ROI in the nucleus (mitochondria free area) and the ratio was plotted over-time, or basal values were given.

### 2-NBDG based glucose uptake measurements

Glucose uptake was assessed using 2-(N-(7-Nitrobenz-2-oxa-1,3-diazol-4-yl)Amino)-2-Deoxyglucose (2-NBDG) (Biomol GmbH). Therefore, cells were seeded on 30 mm glass coverslips (Marienfeld GmbH) in 6-well plates (Corning) and cultivated over-night at 37°C and 5% CO_2_ in a humidified incubator. The next day, cells were taken from the incubator, cell culture medium was replaced for cell equilibration buffer and cells were equilibrated for 30 minutes at ambient environment. Subsequently, cells were washed 3x with glucose free physiologic buffer, and glucose free physiologic buffer containing 100 µM 2-NBDG, with or without paxilline and 200 nM FCCP or an equivalent amount of DMSO, was added to the cells, followed by incubation at 37°C for 30 minutes. Next, cells were washed 3x with glucose free physiologic buffer to remove any residual 2-NBDG and were analyzed by fluorescence microscopy using 485/20 excitation light. Emission was captured at roughly 525 nm (Zeiss Axiovert 200m).

### Mitochondrial single-channel patch-clamp measurements

For mitoplast electrophysiology, mitochondria were isolated from BCC grown to confluency (>90%). Adherent cells were washed 2x with PBS, scraped from the dish, collected in a tube and centrifuged at 400xg for 5 minutes. Subsequently, the cell pellet was resuspended in preparation solution, followed by homogenization using a glass-glass homogenizer. Next, the homogenate was centrifuged at 9,200xg for 10 minutes. The resulting pellet was resuspended in preparation solution and centrifuged at 780xg for 10 minutes. The supernatant was collected, followed by centrifugation at 9,200xg for 10 minutes and resuspension of the mitochondrial fraction in the mitochondrial storage buffer. All procedures were performed at 4°C.

An osmotic swelling procedure was used for the preparation of mitoplasts from the isolated mitochondria. Therefore, mitochondria were added to the hypotonic buffer for ∼1 minute to induce swelling and breakage of the outer mitochondrial membrane. Subsequently, isotonicity of the solution was restored by the addition of hypertonic buffer at a dilution of 1:5.

Patch-clamp experiments on mitoplasts were performed as previously described (Bednarczyk et al., 2013). The experiments were carried out in mitoplast-attached single-channel mode using borosilicate glass pipettes with a mean resistance of 10-15 MΩ. The patch-clamp glass pipette was filled with mitochondrial storage buffer. This isotonic solution was used as a control solution for all experiments. The size of the pipettes and the formation of the gigaseal were monitored by measuring electrical resistance. Connections were made with Ag/AgCl electrodes and an agar salt bridge (3 M KCl) for the ground electrode. The current was recorded using an Axopatch 200B patch-clamp amplifier (Molecular Devices, California, USA). To apply substances, a self-made perfusion system containing a holder with a glass pipe, a peristaltic pump, and Teflon tubing was used. All channel modulators were added as dilutions in the isotonic solution containing 100 μM CaCl_2_.

The presented single-channel current-time recordings are representative for the most frequently observed conductances under the given conditions. The conductance was calculated from the current-voltage relationship. The probability of channel openings and the current amplitude were determined using the Single-Channel Search mode of the Clampfit 10.7 software (Molecular Devices).

### Western blotting

Mitochondria for western blotting were prepared as described earlier(Pallotti and Lenaz, 2007) with some modifications. Cells were washed and collected in PBS and centrifuged at 500xg for 10 min. The pellet was frozen in liquid nitrogen and stored at -80°C. The next day, pellet was resuspended in ice-cold isolation buffer containing (in mM): 210 mannitol, 70 sucrose, 1 PMSF, 5 HEPES and bovine serum albumin (2.5 mg mL^-1^), pH 7.2. Digitonin was added to a final concentration of 0.02–0.04% for additional membrane permeabilization. After 1 min of incubation, digitonin was diluted with isolation buffer and the cells were centrifuged at 3,000xg for 5 min at 4°C. The pellet was resuspended in ice-cold isolation buffer. The cells were homogenized using a glass/glass homogenizer and centrifuged at 1,000xg for 5 min at 4°C followed by another homogenization. The homogenates were centrifuged at 1,000xg for 5 min at 4°C to remove cell remnants. The supernatants were collected and centrifuged for 60 min at 10,000xg, at 4°C. Next, the pellet was resuspended in the isolation buffer without BSA and centrifuged at 10,000xg for 30 min at 4°C, followed by resuspension in isolation buffer without BSA and centrifugation at 10,000xg for 15 min at 4°C. This step was repeated once more. The pellet containing crude mitochondria was resuspended in 0.8 mL of 15% Percoll (in isolation buffer without BSA) and layered on top of a Percoll step gradient (23% and 40% Percoll layers). The suspension was centrifuged at 30,000xg for 30 min at 4°C. Mitochondrial fraction located between the 23% and 40% Percoll layer was collected, diluted with isolation buffer without BSA and centrifuged at 10,000xg for 15 min at 4°C. The final mitochondrial pellet was resuspended in storage buffer containing 500 mM sucrose and 5 mM HEPES, pH 7.2. Each mitochondrial isolation was performed using between 15-30 x 10^6^ cells.

A given amount of sample solubilized in Laemmli buffer (Bio-Rad) was separated by 10% Tris-tricine-SDS-PAGE and transferred onto polyvinylidene difluoride (PVDF) membranes (Bio-Rad). After protein transfer, the membranes were blocked with 10% nonfat dry milk solution in Tris-buffered saline with 0.2% Tween 20 and exposed to one of the following antibodies: anti-BKα antibody (NeuroMabs, USA, clone L6/60, diluted 1:200), anti-COXIV (Cell Signaling Technology, Leiden, Netherlands, no. 4844, 1:1,000), or anti-alpha 1 sodium potassium ATPase (Abcam, Berlin, Germany, no. ab7671, 1:1,000). This was followed by incubation with a secondary anti-rabbit (Thermo Fisher Scientific, no. 31460) or anti-mouse antibody (Thermo Fisher Scientific, no. SA1-100) coupled to horseradish peroxidase. The blots were developed using enhanced chemiluminescence solution (GE Healthcare). To estimate the molecular weight of the analyzed proteins PageRuler Prestained Protein Ladder (Thermo Fisher Scientific) was used.

### qPCR analysis

mRNA was isolated from 6-well plates showing a cell confluency of ∼90% using the Monarch Total RNA Miniprep kit (NEB). siRNA transfection was performed 48 hours prior to RNA isolation using the Lipofectamine 2000 transfection reagent (Thermo Fisher Scientific) according to manufacturer’s instructions. qPCR reactions were performed using the GoTaq 1-Step RT-qPCR System (Promega GmbH, Walldorf, Germany). 100 ng of isolated mRNA were used per reaction. Primers were designed to span exon – exon junctions and to recognize both, human and murine BK_Ca_ sequences. Different primer pairs were used for human and murine b-tubulin. Primer and siRNA sequences are listed in **Supplementary Table 1** and **Supplementary Table 2**. Primers were purchased from Thermo Fisher Scientific, siRNAs were purchased from Microsynth AG. qPCR reactions were run in a CFX connect real-time PCR instrument (Bio-Rad Laboratories GmbH, Feldkirchen, Germany). Sample (Ct) values were normalized to Ct’s of housekeeper gene (b-Tubulin) and calculated as 2^-Ct^ ^(normalized)^.

### Proliferation assays

Proliferation assays were performed using 3-(4,5-dimethylthiazol-2-yl)-2,5-diphenyltetrazolium bromide (MTT) (Thermo Fisher Scientific) assay. For the assay, 2.500 MMTV-PyMT WT and BK-KO cells or 7.500 MDA-MB-453 cells were seeded per well in flat bottom 96-well cell culture plates (Corning) in their respective cell culture medium. Per run, condition, and time-point, 7 technical replicates were performed. One well served as a blank per time-point and condition and did not contain cells. The next day, medium was exchanged for fresh cell culture medium either containing 5 µM paxilline, 30 nM iberiotoxin or an equivalent amount of DMSO as a control and the MTT assay was started for time-point 24 hours directly thereafter. For each time-point (24, 48, 72 and 96 hours), MTT reagent was added with a medium change (10 µL + 100 µL / well) at a final concentration of 455 µg mL^-1^ and cells were incubated for 4 hours. Subsequently, 85 µL of medium were removed from each well, 85 µL of DMSO (Carl Roth GmbH) was added and wells were mixed by pipetting thoroughly. 85 µL of this mixture were transferred to a new 96-well plate (Corning) and absorbance was measured at 540 nm using a TECAN microplate reader (TECAN Group Ltd.). Absorbances at the different time-points (48, 72 and 96 hours) were normalized to the absorbances after 24 hours.

For KI-67 based assessment of cell proliferation rates, 20,000 MMTV-PyMT WT and BK-KO cells and 30,000 MDA-MB-453 cells were seeded in µ-slide 8 wells (ibidi GmbH). The next day, cells were washed 1x with PBS, and serum free cultivation medium was added. Cells were further cultivated for 48 hours, prior to addition of DMSO (ctrl) 5 µM paxilline or 30 nM iberiotoxin. After additional 24 hours, serum containing medium supplemented with DMSO, paxilline or iberiotoxin was re-added, and cell growth was allowed for another 48 hours. Finally, cells were washed 2x with PBS and fixed with ice cold methanol/aceton (50:50 v/v) for 15 minutes at -20°C. Subsequently, cells were washed thrice with PBS, and incubated in PBS containing 5% bovine serum albumin (BSA, Carl Roth GmbH) for 1 hours. Subsequently, anti Ki-67 rabbit monoclonal antibody (D3B5, Cell Signaling Technology) in 5% BSA-PBS was added at a dilution of 1:500. Wells without primary antibody served as a negative control. Slides were incubated over-night at 4°C. The next day, antibody solution was removed, cells were washed 3x with PBS, and subsequently incubated with Goat anti-Rabbit IgG (H+L) Cross-Adsorbed Secondary Antibody, Alexa Fluor 488 (Thermo Fisher Scientific) for 2 hours at room temperature, followed by washing 2x with PBS. During a 3^rd^ washing step, Hoechst 33342 (Thermo Fisher Scientific) at a dilution of 1:2000 was added, cells were incubated for 10 minutes at room temperature and washed an additional time with PBS. Ultimately, cells were mounted in PermaFluor Mountant (Microm International GmbH, Dreieich, Germany), kept over-night at 4°C, and imaged using the Zeiss Axio Observer Z1 wide-field microscope. AlexaFluor 488 and Hoechst 33342 were illuminated using 473/10 and 380x filters, respectively. 2 images were captured per replicate. Image analysis was performed using ImageJ. Threshold positive area of Ki-67 (AlexaFluor488) was expressed as fraction of the threshold positive area of Hoechst 33342 per image, and the average of the 2 images per well was calculated.

### Colony formation assays

For colony formation assays, cell suspensions containing 400 cells mL^-1^ of MMTV-PyMT WT or BK-KO cells were prepared. 1 mL of this suspension was spread drop-per-drop per well of a 6-well cell culture plate (Corning) already containing 1 mL of cell culture medium using 1 mL syringes and 23G needles. Culture plates were gently moved left/right and back/forth to equally distribute the cells across the well, plates were kept at room temperature for 20 minutes to ensure uniform cell settling, and cells were subsequently cultured over-night at 37°C and 5% CO_2_ in a humidified incubator. The next day, medium was exchanged for 2 mL of fresh cell culture medium, and cells were kept in the presence or absence of O_2_ at 37°C and 5% CO_2_ in humidified incubators for 7 days. After 7 days, cells were washed carefully 2x with PBS and fixed for 30 minutes on ice using 2% PFA in PBS. Meanwhile, a solution of crystal violet (0.01% w/v) was prepared in ddH_2_O, cells were washed 2x with PBS and incubated in the crystal violet solution (Sigma Aldrich Chemie GmbH) for 60 minutes after fixation, followed by excessive washing with ddH_2_O until a clear background was obtained. Plates were dried at room temperature for at least 12 hours before images were captured using an Amersham Imager 600 (GE Healthcare UK Limited, Buckinghamshire, UK). Analysis was performed using ImageJ.

### Breast cancer patient study

A subset of 551 primary tumor specimens with available archival tumor blocks were obtained from a prospective, observational multicenter adjuvant endocrine treatment study of 1286 post-menopausal HR-positive breast cancer patients recruited between 2005 and 2011 (German Clinical Trial Register DRKS 00000605, “IKP211” study). Study inclusion and exclusion criteria have been previously described (Schroth et al., 2020). Patients had received standard endocrine treatment, i.e. tamoxifen, aromatase inhibitors, or switch regimens between both. Ethics approval was obtained from the Medical Faculty of the University of Tuebingen and the local ethics committees of all participating centers in Germany. Informed patient consent was obtained from all participants as required by institutional review boards and research ethics committees. All patient data were de-identified prior inclusion in this study.

### Nanostring nCounter gene expression analysis

Gene expression analysis was performed according to manufacturer’s protocol. In brief, total RNA from human primary tumors of the IKP211 study (Quick-DNA/RNA FFPE, ZymoResearch, Freiburg, Germany) was extracted from 10 µm sections of FFPE tissues with at least 20% tumor cell content. Samples were enriched for tumor tissue by microdissection. A total of 250 ng RNA was subsequently hybridized with target-specific capture and color-coded reporter probe sets in a pre-warmed thermal cycler at 65°C for 20 h. In the post-hybridization process, the total volume was increased to 32 µl with RNase-free water. Fluorescence count measurements were immediately conducted in the Nanostring nCounter System. Data were analyzed using nSolver 4.0 and normalized to housekeeping genes. ABCF1, NRDE2, POLR2A, PUM1 and SF3A1 served as housekeepers. Probes used for Nanostring nCounter gene expression analysis are listed in **Supplementary Table 3**.

### Statistical analysis

Statistical analysis was performed using Prism8 software (GraphPad Software, Boston, USA). All data were tested for normal distribution using D’Agostino and Pearson omnibus normality test. A two-tailed Unpaired t-test or Welch’s t-test were used for statistical comparison of normally distributed data, depending on whether the variances of the populations were comparable or significantly different as tested by an F-test. A two-tailed Mann-Whitney test was used for pairwise comparison of non-normally distributed data. Comparison of >2 data sets was done either using One-way ANOVA followed by Tukey’s multiple comparison (MC) test or Brown-Forsythe and Welch ANOVA test followed by Games-Howell’s MC test for normally distributed data, depending on whether the variances were comparable or significantly different. A Kruskal-Wallis test followed by Dunn’s MC test was performed if data were not normally distributes. The statistical tests used are indicated in the figure legends. p-values of ≤0.05 were considered as significant, where * or #p≤0.05, ** or †p≤0.01 and *** or ‡p≤0.001. No priory sample size estimation was performed.

## Acknowledgements

This work was funded by the Deutsche Forschungsgemeinschaft (DFG) with individual grants to RL and LM. RL is a member of the GRK2381: “cGMP: From Bedside to Bench”, DFG grant number 335549539. SM, WS, MS and RL acknowledge financial support from the ICEPHA Graduate Program “Membrane-associated Drug Targets in Personalized Cancer Medicine”. HB is a fellow of the Austrian Science Fund (FWF) funded Erwin-Schrödinger-Program, project number J-4457. SB acknowledges financial support from the Fritz Thyssen Stiftung. The authors thank Clement Kabagema-Bilan, Michael Glaser and Antoni Wrzosek for excellent technical support and Peter Ruth for valuable discussions. This work was partially supported by the Nencki Institute of Experimental Biology, the Polish National Science Centre grant no. 2019/34/A/NZ1/00352 (AS), and by the FWF, project number I-3716 (RM).

## Author contributions

HB and RL initiated and designed this study; HB, SM, PK, BK, SB, JJ, KS, YZ, WS and PB performed experiments and were involved in data curation; HB, SM, PK, BK, SB, JJ, KS and WS assisted in formal analysis; HB, SM, SB and PB visualized data; HB, SB, WS, LM, MS, PB, AS and RL acquired financial support; HB, SM, PK, BK, SB, KS, WS, ML and PB designed the methodology; HB and RL performed project administration; PK, WS, LM, MS, ALB, RM, ML, PB, AS and RL provided resources; RL supervised the project and the personnel; HB and RL wrote the original draft. All authors critically reviewed the manuscript and stated comments.

## Data availability

Raw data for the graphs can be found in source data files.

