## Supplementary Information for "mitoBK_Ca_ is functionally expressed in murine and human breast cancer cells and potentially contributes to metabolic reprogramming"

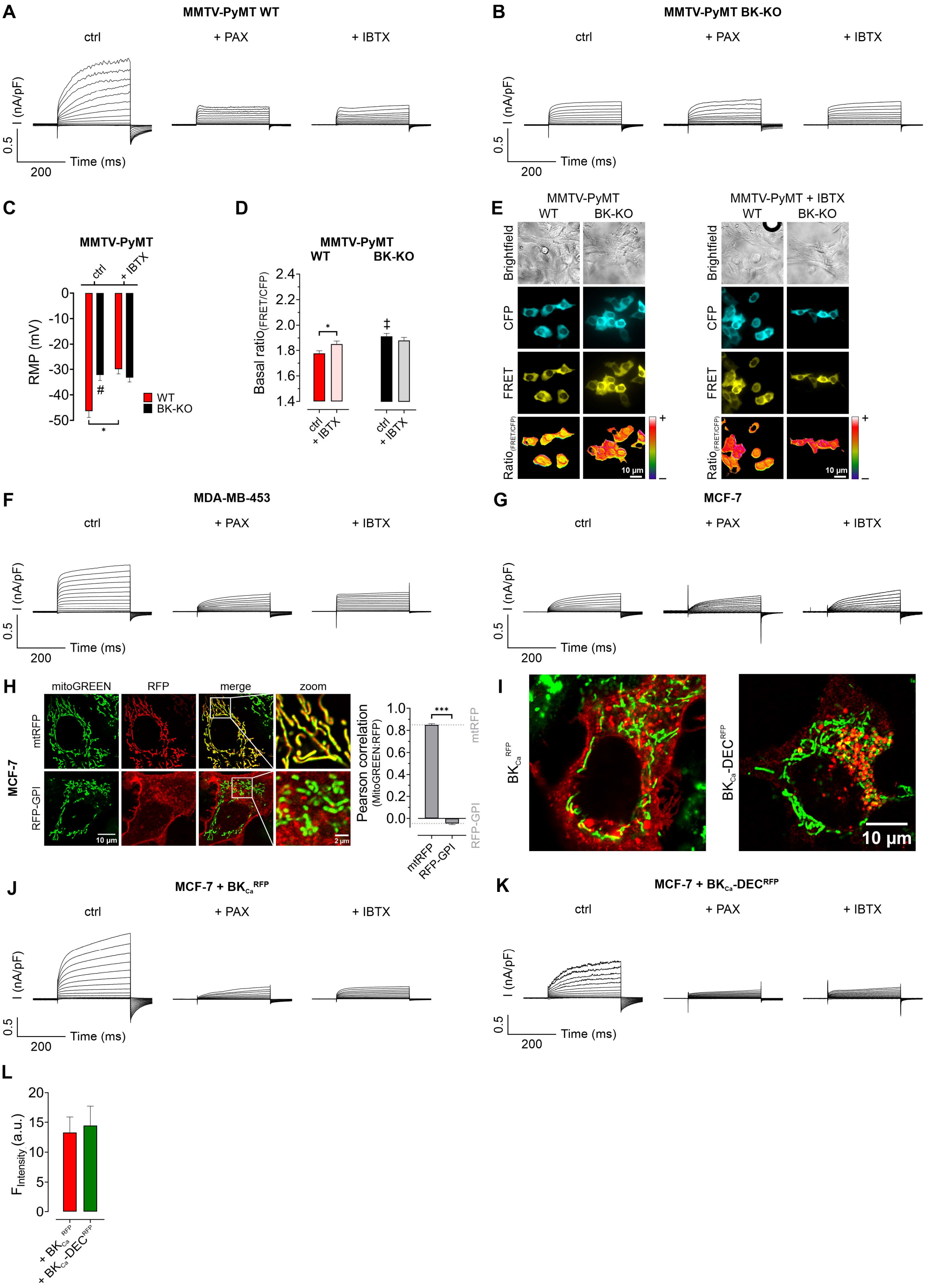

**Supplementary Figure 1: Representative whole-cell patch-clamp traces and colocalization analysis in BCCs.** (**A, B**) Representative whole-cell patch-clamp traces of MMTV-PyMT WT (**A**) and MMTV-PyMT BK-KO cells (**B**), either under control conditions (ctrl), or in the presence of 5 µM paxilline (+ PAX) or 30 nM iberiotoxin (+ IBTX), respectively, as indicated in the panels. (**C**) Resting membrane potential (RMP) ± SEM of MMTV-PyMT WT (red) and BK-KO cells (black) in mV as measured using current clamp mode during patch-clamp experiments. Cells were either analyzed under control conditions (ctrl, left bars), or in the presence of 30 nM iberiotoxin (+ IBTX, right bars). n (cells) = 9 for WT ctrl and WT + IBTX, 8 for BK-KO ctrl, 10 for BK-KO + IBTX. *p≤0.05, #p≤0.05 compared to respective WT condition. Statistical analysis was performed using Kruskal-Wallis test followed by Dunn's multiple comparison test. (**D**) Basal FRET-ratio values ± SEM of MMTV-PyMT WT (left panel) and MMTV-PyMT BK-KO cells (right panel) expressing NES lc-LysM GEPII 1.0, a cytosolic $K^+$ sensor. Experiments were either performed under control conditions (ctrl) or in the presence of 30 nM iberiotoxin (+ IBTX). N (independent experiments) / n (cells analyzed) = 4/62 WT ctrl, 4/50 WT + IBTX, 4/35 BK-KO ctrl, 4/36 BK-KO + IBTX. *p≤0.05. ‡p≤0.001 compared to respective WT condition, Unpaired t-test (WT) or Mann-Whitney test (BK-KO and WT ctrl vs. BK-KO ctrl). (**E**) Representative images of MMTV-PyMT WT and MMTV-PyMT BK-KO cells, either under control conditions (left panel) or in the presence of 30 nM iberiotoxin (+ IBTX, right panel). Brightfield images (top row), cyan fluorescence (second row), FRET (third row) and pseudocolored ratio images (fourth row) are demonstrated. (**F, G**) Representative whole-cell patch-clamp traces of MDA-MB-453 cells (**F**) and MCF-7 cells (**G**), either under control conditions (ctrl), or in the presence of 5 µM paxilline (+ PAX) or 30 nM iberiotoxin (+ IBTX), respectively, as indicated in the panels. (**H**) Representative images (left) of MCF-7 cells either expressing a mitochondrial targeted red fluorescent protein (mtRFP, upper images, second column) or a red fluorescent protein fused to a glycosylphosphatidylinositol (GPI)-anchor (RFP-GPI, lower images, second column). Cells were additionally stained with MitoGREEN for visualization of mitochondria (first column). Merge of the channels and a zoom are demonstrated. Right panel shows average Pearson correlation ± SEM of MitoGREEN and RFP of mtRFP (left bar) or RFP-GPI (right bar). Grey dashed lines indicate average colocalization scores of MitoGREEN and RFP of mtRFP and RFP-GPI, which is also shown in Figure 1G. n (cells) = 18 for mtRFP and 16 for RFP-GPI. ***p≤0.001 Unpaired t-test. (**I**) Representative images of MCF-7 cells either expressing $BK_{Ca}^{RFP}$ (left image) or $BK_{Ca}$-$DEC^{RFP}$ (right image) additionally stained with MitoGREEN to visualize mitochondria. Images represent larger versions of the exact same images shown in Figure 1G. (**J, K**) Representative whole-cell patch-clamp traces of MCF-7 cells either expressing $BK_{Ca}^{RFP}$ (**J**) or $BK_{Ca}$-$DEC^{RFP}$ (**K**) under control conditions (ctrl, left panels), or in the presence of 5 µM paxilline (+ PAX, middle panels) or 30 nM iberiotoxin (+ IBTX, right panels). (**L**) Global cellular RFP-fluorescence intensities of MCF-7 cells used for patch-clamp experiments. Cells either expressed $BK_{Ca}^{RFP}$ (red bar) or $BK_{Ca}$-$DEC^{RFP}$ (green bar). Data represents average ± SEM of 18 cells for both conditions.

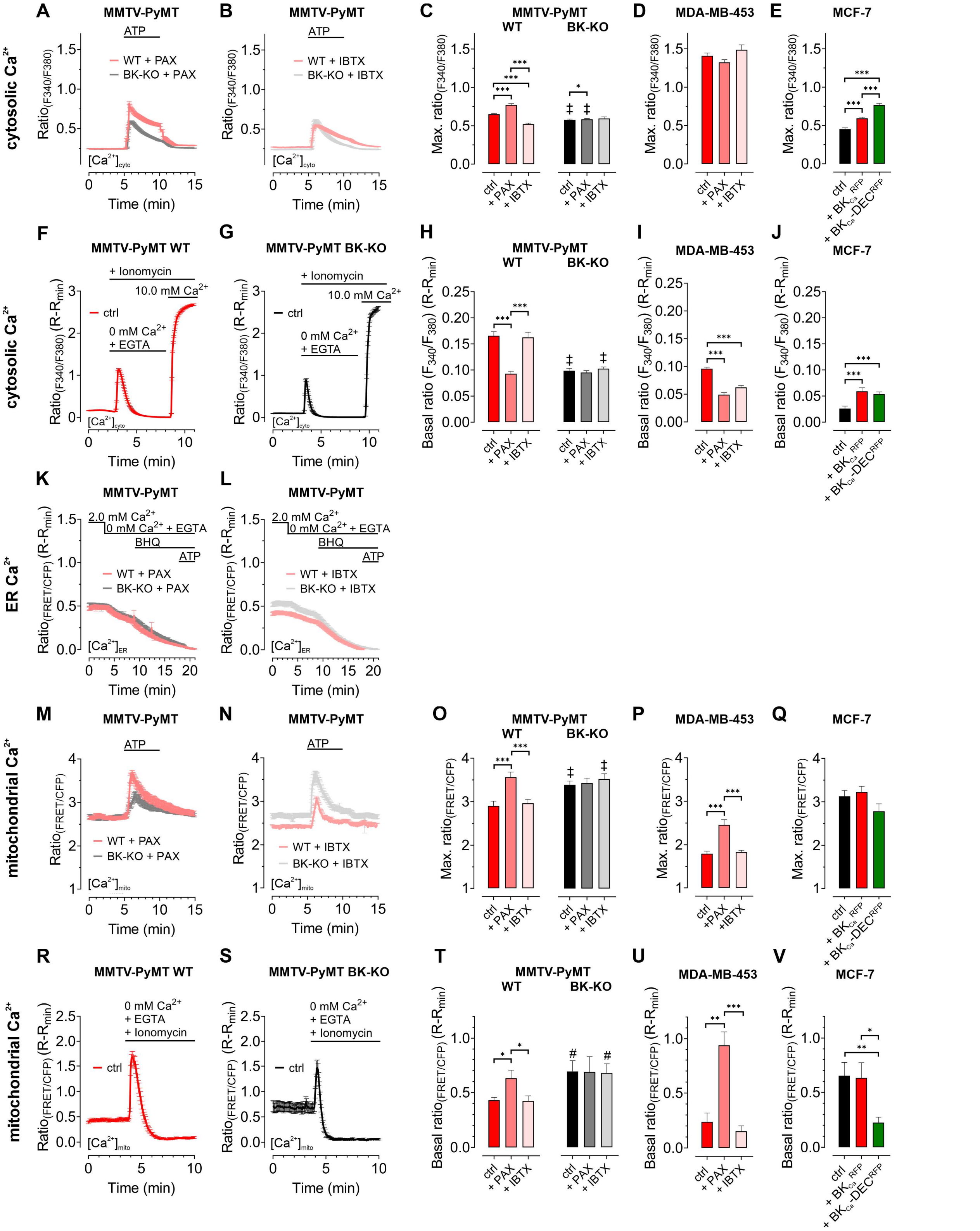


**Supplementary Figure 2: Cytosolic-, ER-, and mitochondrial $Ca^{2+}$ homeostasis is altered by functional $BK_{Ca}$ expression in BCCs.** (**A, B**) Fura-2 ratio signals ($F_{340}/F_{380}$) of MMTV-PyMT WT or BK-KO cells over-time in response to cell stimulation with 100 µM ATP. Experiments were either performed in the presence of 5 µM paxilline (**A**) or 30 nM iberiotoxin (**B**). (**C – E**) Maximal fura-2 ratio signals of MMTV-PyMT WT and BK-KO (**C**), MDA-MB-453 (**D**) or MCF-7 cells (**E**) upon stimulation with 100 µM ATP, under control conditions, in the presence of 5 µM paxilline or 30 nM iberiotoxin (**C, D**), or upon expression of $BK_{Ca}^{RFP}$ or $BK_{Ca}$-$DEC^{RFP}$ (**E**). (**F, G**) Fura-2 ratio signals ± SEM over-time of MMTV-PyMT WT and BK-KO cells. At time points indicated in the panels, cells were treated with 5 µM ionomycin for

$Ca^{2+}$ permeabilization, $Ca^{2+}$ was removed and chelated using EGTA, or 10.0 mM of $Ca^{2+}$ were re-added for Fura-2 saturation. (**H – J**) Basal Fura-2 ratio signals ($R-R_{min}$) ± SEM of MMTV-PyMT WT and BK-KO cells (**H**), MDA-MB-453 (**I**) and MCF-7 cells (**J**) received from experiments as demonstrated in (**F**) and (**G**). (**K, L**) FRET-ratio signals over-time of MMTV-PyMT WT and BK-KO cells expressing D1ER, a FRET-based ER $Ca^{2+}$ sensor. Throughout the experiments, either 5 µM paxilline (**F**) or 30 nM iberiotoxin (**G**) were present. (**M, N**) $[Ca^{2+}]_{mito}$ over-time of MMTV-PyMT WT or BK-KO cells in response to cell stimulation with 100 µM ATP. $[Ca^{2+}]_{mito}$ was assessed using 4mtD3cpV, a FRET-based $Ca^{2+}$ indicator targeted to the mitochondrial matrix. Experiments were either performed in the presence of 5 µM paxilline (**M**) or 30 nM iberiotoxin (**N**). (**O – Q**) Maximal FRET-ratio (FRET/CFP) signals received upon stimulation of MMTV-PyMT WT or BK-KO (**O**), MDA-MB-453 (**P**) or MCF-7 cells (**Q**) with 100 µM ATP. Experiments were either performed under control conditions, in the presence of 5 µM paxilline or 30 nM iberiotoxin (**O, P**), or upon expression of $BK_{Ca}^{RFP}$ or $BK_{Ca}$-$DEC^{RFP}$ (**Q**). (**R, S**) FRET-ratio signals over-time of MMTV-PyMT WT and BK-KO cells expressing 4mtD3cpV, a FRET-based mitochondrial $Ca^{2+}$ sensor. At time point indicated in the panels, cells were treated with 5 µM ionomycin for $Ca^{2+}$ permeabilization, and $Ca^{2+}$ was removed and chelated using EGTA. (**T – V**) Basal FRET-ratio signals ($R-R_{min}$) of MMTV-PyMT WT and BK-KO cells (**T**), MDA-MB-453 (**U**) and MCF-7 cells expressing 4mtD3cpV (**V**) received from experiments as demonstrated in (**R**) and (**S**). All data represent average ± SEM. N (independent experiments) / n (cells analyzed) = **A:** 6/300 WT + PAX, 6/300 BK-KO + PAX, **B:** 5/318 WT + IBTX, 5/304 BK-KO + IBTX, **C:** 17/784 WT ctrl, 18/857 BK-KO ctrl, 6/300 WT + PAX, 6/300 BK-KO + PAX, 5/318 WT + IBTX, 5/304 BK-KO + IBTX, **D:** 4/151 ctrl, 4/132 + PAX, 4/87 + IBTX, **E:** 5/111 ctrl, 5/117 + $BK_{Ca}^{RFP}$, 5/91 + $BK_{Ca}$-$DEC^{RFP}$, **F:** 3/109 WT ctrl, **G:** 3/93 BK-KO ctrl, **H:** 3/109 WT ctrl, 3/110 WT + PAX, 3/123 WT + IBTX, 3/93 BK-KO ctrl, 3/94 BK-KO + PAX, 3/111 BK-KO + IBTX, **I:** 3/106 ctrl, 3/107 + PAX, 3/109 + IBTX, **J:** 4/53 ctrl, 5/34 + $BK_{Ca}^{RFP}$, 5/36 + $BK_{Ca}$-$DEC^{RFP}$, **K:** 8/71 WT + PAX, 8/92 BK-KO + PAX, **L:** 6/102 WT + IBTX, 6/86 BK-KO + IBTX, **M:** 6/46 WT + PAX, 6/58 BK-KO + PAX, **N:** 5/59 WT + IBTX, 4/43 BK-KO + IBTX, **O:** 11/47 WT ctrl, , 6/46 WT + PAX, 5/59 WT + IBTX, 12/86 BK-KO ctrl, 6/58 BK-KO + PAX, 4/43 BK-KO + IBTX, **P:** 8/33 ctrl, 8/28 + PAX, 5/22 + IBTX, **Q:** 5/28 ctrl, 4/27 + $BK_{Ca}^{RFP}$, 4/24 + $BK_{Ca}$-$DEC^{RFP}$, **R:** 3/19 WT ctrl, **S:** 3/12 BK-KO ctrl, **T:** 3/19 WT ctrl, 3/23 WT + PAX, 3/22 WT + IBTX, 3/12 BK-KO ctrl, 3/17 BK-KO + PAX, 3/14 BK-KO + IBTX, **U:** 6/16 ctrl, 6/19 + PAX, 6/19 + IBTX, **V:** 5/21 ctrl, 4/10 + $BK_{Ca}^{RFP}$, 4/11 + $BK_{Ca}$-$DEC^{RFP}$**.** *p≤0.05, **p≤0.01, ***p≤0.001, Kruskal-Wallis test followed by Dunn's MC test (**C, H, I, J, T, U**), One-Way ANOVA test followed by Tukey's MC test (**E, O**) or Brown-Forsythe and Welch ANOVA test followed by Games-Howell's MC test (**P, V**). #p≤0.05, ‡p≤0.001 compared to respective WT condition, Mann-Whitney test (**H, O,** ctrl in **C,** + IBTX in **T**), Welch's t-test (+ PAX in **C,** ctrl in **T**). Unpaired t-test (+ IBTX in **O**).

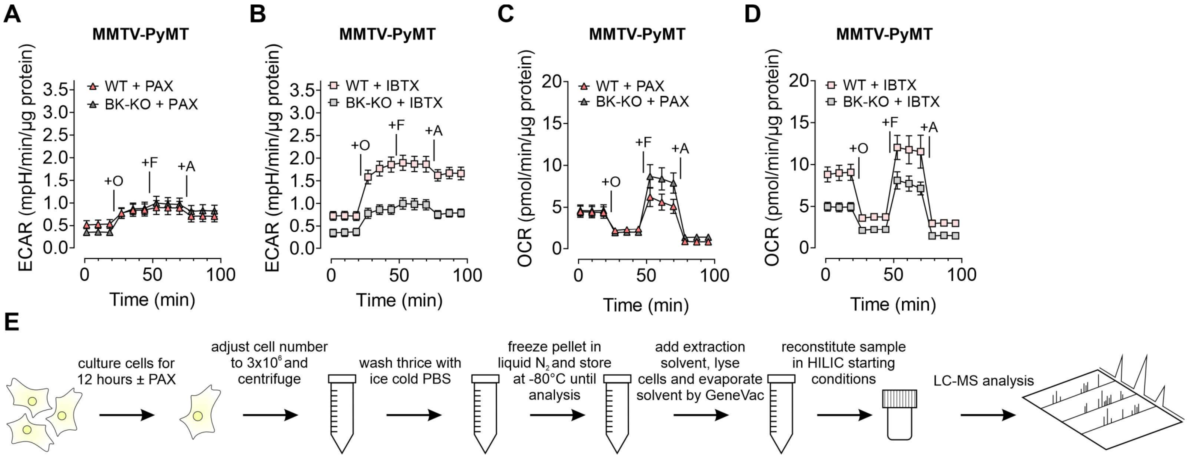


**Supplementary Figure 3: Paxilline and iberiotoxin differentially modulate ECAR and OCR in MMTV-PyMT WT and BK-KO cells.** (**A, B**) Average ECAR over-time ± SEM of MMTV-PyMT WT (bright red and salmon) and BK-KO cells (dark and bright grey) in response to administration of Oligomycin-A (+O), FCCP (+F) and Antimycin-A (+A) at time points indicated in the panel. Experiments were either performed in the presence of 5 µM paxilline (**A**) or 30 nM iberiotoxin (**B**). N = 3 for all. (**C, D**) Average OCR over-time ± SEM of MMTV-PyMT WT (bright red and salmon) and BK-KO cells (dark and bright grey) in response to administration of Oligomycin-A (+O), FCCP (+F) and Antimycin-A (+A) at time points indicated in the panel. Experiments were either performed in the presence of paxilline (**C**) or iberiotoxin (**D**). N = 3 for all. (**E**) Schematic representation of the workflow for LC-MS-based metabolomics. HILIC: Hydrophilic interaction liquid chromatography.

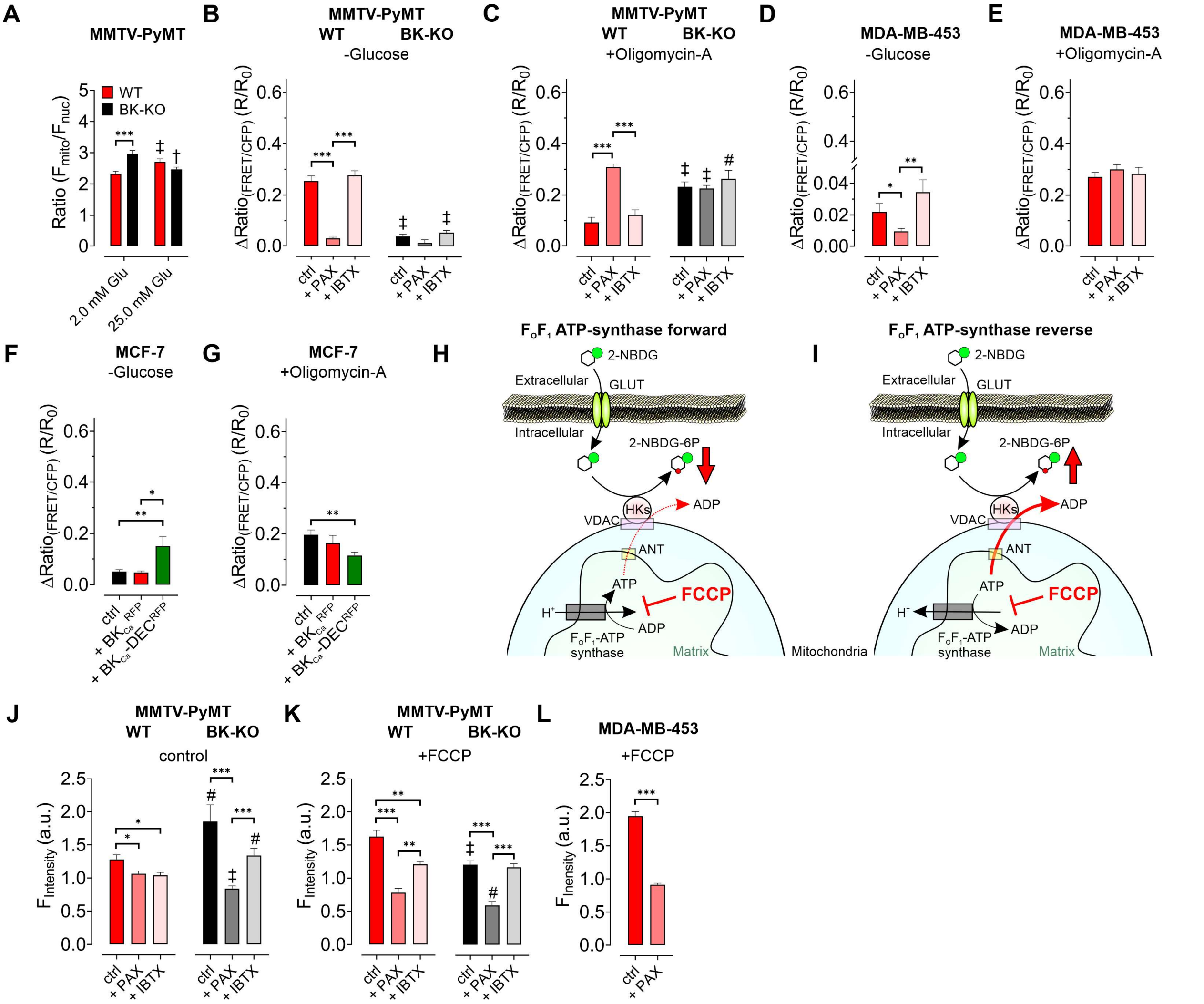


**Supplementary Figure 4: $BK_{Ca}$ modulates cellular substrate dependency for maintaining $[ATP]_{mito}$ and reverses $F_OF_1$ ATP-synthase.** (**A**) Average fluorescence ratios ($F_{mito}/F_{nuc}$) of MMTV-PyMT WT and BK-KO cells, either in the presence of 2.0 mM or 25.0 mM extracellular glucose. N (independent experiments) / n (cells analyzed) = 6/171 WT 2.0 mM Glu, 6/180 WT 25.0 mM Glu, 5/133 BK-KO 2.0 mM Glu, 6/163 BK-KO 25.0 mM Glu. ***$p \leq 0.001$, Mann-Whitney test. †$p \leq 0.01$, ‡$p \leq 0.001$ compared to 2.0 mM glucose condition of the respective cell type, Mann-Whitney test. (**B – E**) Average changes in FRET-ratio signals ± SEM induced either upon extracellular glucose removal (**B, D**) or upon administration of Oligomycin-A (**C, E**) to MMTV-PyMT WT, BK-KO (**B, C**) or MDA-MB-453 cells (**D, E**) expressing mtAT1.03, a FRET-based ATP sensor targeted to the mitochondrial matrix. Experiments were either performed under control conditions or in the presence of 5 µM paxilline or 30 nM iberiotoxin. N/n = **B:** 8/55 WT ctrl, 6/45 WT + PAX, 7/27 WT + IBTX, 8/65 BK-KO ctrl, 6/57 BK-KO + PAX, 7/28 BK-KO + IBTX, **C:** 11/52 WT ctrl, 7/53 WT + PAX, 7/34 WT + IBTX, 8/87 BK-KO ctrl, 6/35 BK-KO + PAX, 8/45 BK-KO + IBTX, **D:** 5/14 ctrl, 3/13 + PAX and 5/13 + IBTX, **E:** 5/33 ctrl, 3/21 + PAX, 8/27 + IBTX. *$p \leq 0.05$, **$p \leq 0.01$, ***$p \leq 0.001$, Kruskal-Wallis test followed by Dunn's MC test (**B, C, D**). #$p \leq 0.05$, ‡$p \leq 0.001$ compared to respective WT condition, Mann-Whitney test (ctrl in **B**, all in **C**), or Welchs t-test (+ IBTX in **B**). (**F, G**) Average changes in FRET-ratio signals ± SEM induced either upon extracellular glucose removal (**F**) or upon administration of Oligomycin-A (**G**) to MCF-7 cells expressing mtAT1.03, either in combination with a red fluorescent protein as control, or $BK_{Ca}^{RFP}$ or $BK_{Ca}$-$DEC^{RFP}$, respectively *$p \leq 0.05$, **$p \leq 0.01$, Kruskal-Wallis test followed by Dunn's MC test. N/n = **F:** 6/48 ctrl, 5/23 + $BK_{Ca}^{RFP}$, 5/20 + $BK_{Ca}$-$DEC^{RFP}$, **G:** 5/27 ctrl, 5/23 + $BK_{Ca}^{RFP}$, 5/37 + $BK_{Ca}$-$DEC^{RFP}$. (**H, I**) Schematic representation of processes involved in 2-NBDG uptake. 2-NBDG is taken up via glucose transporters (GLUTs, green), and ATP-dependently phosphorylated by hexokinase isoforms (HKs, red) to 2-NBDG-6-phosphate (2-NBDG-6P). Under basal conditions, if $F_OF_1$ ATP-synthase is running in forward mode (**H**), mitochondria contribute to 2-NBDG uptake by ATP generation and delivery to HKs via the adenine nucleotide transporter (ANT, yellow), and the voltage-dependent anion channel (VDAC, violet). Contrary, under basal conditions, if $F_OF_1$ ATP-synthase activity is reversed (**I**) it competes with HKs for ATP. Subsequent inhibition of mitochondria due to their depolarization with FCCP (red) stops ATP-synthase activity. Under these conditions, 2-NBDG uptake will decrease if $F_OF_1$ ATP-synthase operates in forward mode (**H**), but will increase if $F_OF_1$ ATP-synthase shows reversed activity (**I**). (**J – L**) Average fluorescence signal (a.u.) ± SEM of MMTV-PyMT WT and BK-KO cells (**J, K**), or MDA-MB-453 cells (**L**) incubated with 100 μM 2-NBDG at 37°C for 30 minutes either under control conditions (**J**) or in the presence of 200 nM FCCP for mitochondrial depolarization (**K, L**), with or without 5 μM paxilline or 30 nM iberiotoxin. N = 4 for all. *$p \leq 0.05$, ***$p \leq 0.001$, Kruskal-Wallis test followed by Dunn's MC test (**J**, and WT in **K**), One-Way ANOVA test (BK-KO in **K**) or Welch's t-test (**L**). #$p \leq 0.05$, ‡$p \leq 0.001$ compared to respective WT condition, Mann-Whitney test (ctrl and + IBTX in **J**, ctrl in **K**), or Unpaired t-test (+ PAX in **J** and + PAX in **K**).

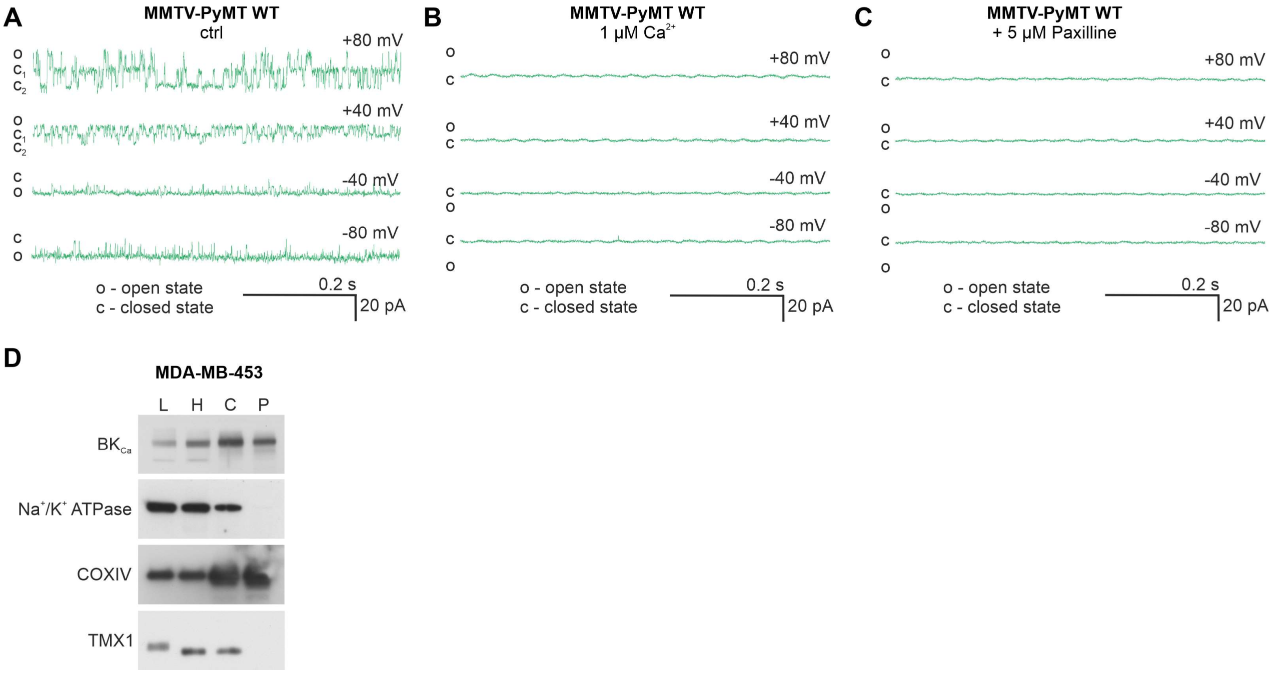


**Supplementary Figure 5: $BK_{Ca}$ is present in the inner mitochondrial membrane of MMTV-PyMT WT and MDA-MB-453 cells.** (**A – C**) Graphs show representative $BK_{Ca}$ single-channel recordings of the inner mitochondrial membrane of mitoplasts isolated from MMTV-PyMT WT cells using a symmetric 150/150 mM isotonic KCl solution, either containing 100 µM $Ca^{2+}$ (**A**), 1 µM $Ca^{2+}$ (**B**), or 5 µM Paxilline in the presence of 100 µM $Ca^{2+}$ (**C**), at voltages ranging from -80 to +80 mV as indicated in the panels. The patch in **a** contained two $BK_{Ca}$ channels. "c" indicates the closed-, "o" the open state of the channel. (**D**) Representative western blot of $BK_{Ca}$ ), $Na^+/K^+$ ATPase as a plasma membrane marker, cytochrome c oxidase subunit IV (COXIV) as a mitochondrial marker and thioredoxin-related transmembrane protein 1 (TMX1), a protein enriched in the ER membrane. Western blot was performed using whole-cell lysates (L), the homogenate (H), crude isolated mitochondria (C) and mitochondria after percoll gradient purification (P) of MDA-MB-453 cells. N = 3.

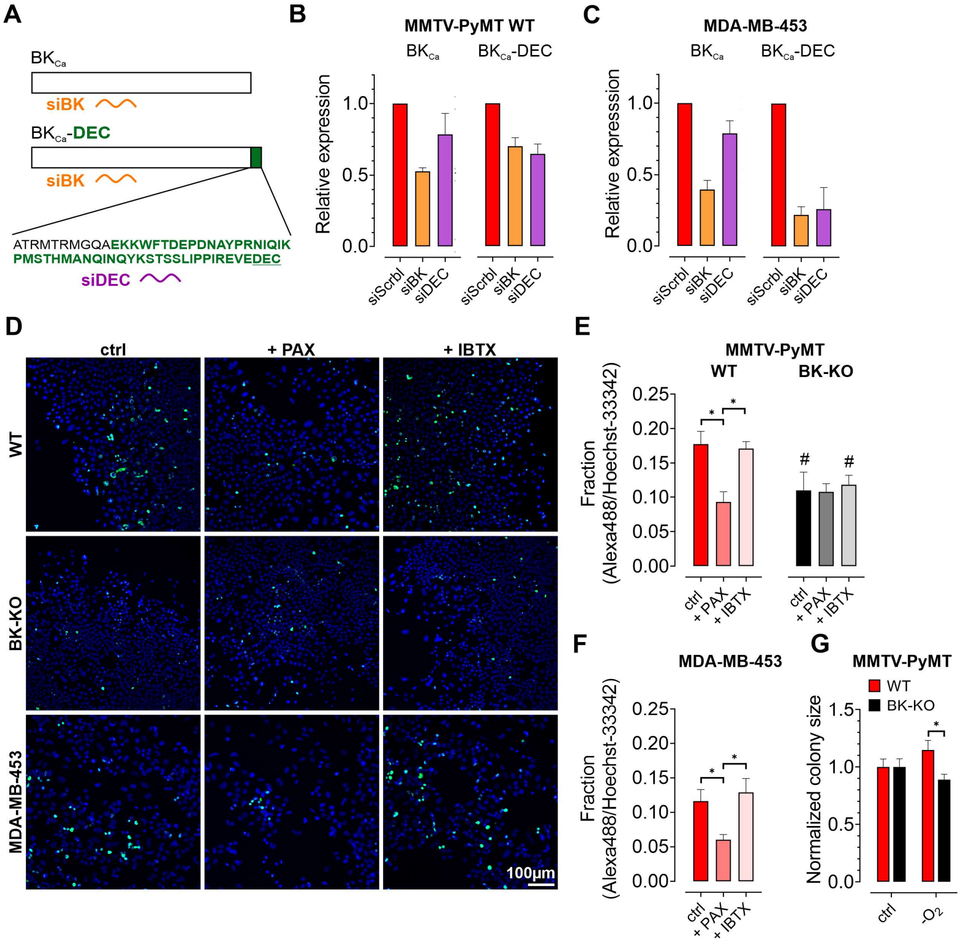


**Supplementary Figure 6: Effects of siRNA treatment on expression of BK$_{Ca}$ and BK$_{Ca}$-DEC, and physiologic consequences of BK$_{Ca}$ inhibition in BCCs.** (**A**) Schematic representation of siRNAs used for subsequent silencing experiments. Either an siRNA targeting all known BK$_{Ca}$ isoforms, referred to as siBK (orange), or an siRNA specifically designed against the DEC exon (green), referred to as siDEC (violet), were used. (**B, C**) Relative mRNA expression levels ± SEM of BK$_{Ca}$ and BK$_{Ca}$-DEC in MMTV-PyMT WT (**B**) and MDA-MB-453 cells (**C**), as analyzed by qPCR. Cells were either treated with a scrambled siRNA as a control (siScrbl), siRNA against all known BK$_{Ca}$ isoforms (siBK) or siRNA specifically targeting BK$_{Ca}$-DEC (siDEC), respectively. N = 4 siBK MMTV-PyMT WT, 3 for all others. (**D-F**) Representative images (**D**) and corresponding statistics (**E, F**) of MMTV-PyMT WT cells (**D**, upper line and **E**, left panel), MMTV-PyMT BK-KO cells (**D**, middle line and **E**, right panel) and MDA-MB-453 cells (**D**, lower line, and **F**) immune-stained for the proliferation marker KI-67 (Alexa488, green), and Hoechst-33342 (blue) for visualizing all nuclei. Experiments were either performed under control conditions (ctrl, left column), or in the presence of 5 µM paxilline (+ PAX, middle column) or iberiotoxin (+ IBTX, right column). *$p \leq 0.05$, Kruskal-Wallis test followed by Dunn's MC test. #$p \leq 0.05$ compared to respective WT condition, Mann-Whitney test. N = 5 for WT ctrl, BK-KO + PAX and MDA-MB-453 + IBTX, N = 6 for all others. (**G**) Normalized colony sizes of colony formation assays performed using MMTV-PyMT WT and BK-KO cells. Cells were cultivated for 7 days either in the presence or absence of $O_2$. *$p \leq 0.05$, Mann-Whitney test. N = 4 independent experiments for all conditions.

**Supplementary Table 1: Probes used for Nanostring nCounter gene expression analysis.**

| Gene | Accession number | Target sequence (5' – 3') |
|---|---|---|
| ABCF1 | NM_001090.2 | TCCCGCCAAGCCATGTTAGAAAATGCATCTGACATCAAGCTGGAGAAGTTCAGCATCTCCGCTCATGGCAAGGAGCTGTTCGTCAATGCAGACCTGTACA |
| NRDE2 | NM_017970.3 | TGGAGGTCCTATGTACAGATTCAGAATAAGTCCCACAGTGCCAGCAAAACCAGGAGATTTTTTGACACAATCACCAGGTCTGCCAAACCCTTGGAGCCTT |
| POLR2A | NM_000937.2 | TTCCAAGAAGCCAAAGACTCCTTCGCTTACTGTCTTCCTGTTGGGCCAGTCCGCTCGAGATGCTGAGAGAGCCAAGGATATTCTGTGCCGTCTGGAGCAT |
| PUM1 | NM_001020658.1 | CTGGGGAACATCAGATCATTCAGTTTCCCAGCCAATCATGGTGCAGAGAAGACCTGGTCAGAGTTTCCATGTGAACAGTGAGGTCAATTCTGTACTGTCC |
| SF3A1 | NM_005877.4 | GATGATGAGGTGTACGCACCAGGTCTGGATATTGAGAGCAGCTTGAAGCAGTTGGCTGAGCGGCGTACTGACATCTTCGGTGTAGAGGAAACAGCCATTG |
| KCNMA1 | NM_001014797.2 | CCGTGCGACAGCCGGGGCCAACGCATGTGGTGGGCTTTCCTGGCCTCCTCCATGGTGACTTTCTTCGGGGGCCTCTTCATCATCTTGCTCTGGCGGACGC |
| KCNMA1-DEC | XM_024447988.2 | AAACAGAATGCAACAAGGATGAATAGAATGGGCCAAGAAAAGAAATGGTTTACAGATGAACCGGATAATGCCTATCCCAGAAACATTCAAATCAAGCCCA |

**Supplementary Table 2: Primers used for qPCR analysis.**

| Primer name | Sequence (5' – 3') | Species | Amplicon size |
|---|---|---|---|
| $BK_{Ca}$-DEC for | CAAACAGAATGCAACAAGGATG | Human / mouse | 124 bp |
| $BK_{Ca}$-DEC rev | GTTAGCCATGTGGGTACTC | Human / mouse | |
| $BK_{Ca}$ for | CGCCTCTTCATGGTCTTC | Human / mouse | 134 bp |
| $BK_{Ca}$ rev | ATGTGCTTTCTTCCACTAAC | Human / mouse | |
| h β-tubulin for | GGCCAGATCTTTAGACCAGAC | Human | 120 bp |
| h β-tubulin rev | CACATCCAGGACAGAATCAAC | Human | |
| m β-tubulin for | AGTGTGGCAACCAGATC | Mouse | 114 bp |
| m β-tubulin rev | AGTAAACGCTGATCCTCTC | Mouse | |

**Supplementary Table 3: siRNAs used for silencing based experiments.**

| siRNA name | Sequence (5’ – 3’) | Targeted Species |
|---|---|---|
| siScrbl | UUCUCCGAACGUGUCACGU-dTdT | Human / mouse |
| siBK | UAGGAAACCGCAAGAAAUA-dTdT | Human / mouse |
| siBK-DEC | CCAGAUCAACCAAUAUAAA-dTdT | Human / mouse |
